# Identification and Characterization of Stem Cells in Mammalian Esophageal Stratified Squamous Epithelia

**DOI:** 10.1101/2021.11.11.468184

**Authors:** Yanan Yang, Guodong Deng, Lili Qiao, Hui Yuan, Xiaohong Yu, Lei Xu, Shih-Hsin Lu, Wei Jiang, Xiying Yu

## Abstract

Somatic stem cells are essential for maintenance of cell proliferation-differentiation homeostasis in organs. Despite the importance, how the esophageal epithelium that executes its self-renewal and maintenance remains elusive. In this study, using 5-bromo-2’-deoxyuridine (BrdU) label-chase in rat and rat esophageal keratinocyte cell line-derived organoids together with genome-wide DNA methylation profiling and single-cell RNA sequencing (scRNA-seq), we identify slow cycling/quiescent stem cell population that contain high levels of hemidesmosome (HD)’s and low levels of Wnt signaling localized spatially and randomly at the basal layer of the esophageal epithelium. Pseudo-time cell trajectory from scRNA-seq indicates that cell fates begin from quiescent basal cells (the stem cells) of the basal layer that produce proliferating and/or differentiating cells in the basal layer, which, in turn, progress into differentiating cells in the suprabasal layer, ultimately transforming into differentiated keratinocytes in the differentiated layer. Perturbations of HD component expressions and/or Wnt signaling reduce stem cell in the basal layer of esophageal keratinocyte organoids, resulting in alterations of organoid formation rate, size, morphogenesis and proliferation-differentiation homeostasis. Furthermore, we show that not only high levels of HDs and low levels of Wnt signaling but also an interplay between HD and Wnt signaling defined stem cells of the basal layer in the esophageal squamous epithelium. Hence, HDs and Wnt signaling are the critical determinants for defining stem cells of the basal layer required for proliferation-differentiation homeostasis and maintenance in the mammalian esophageal squamous epithelium.

## Introduction

The endoderm-derived esophagus in mammals is an important organ of the digestive system between the oropharynx and the stomach for transporting ingested foods. The mammalian esophagus initially originates from the anterior foregut that also gives rise to the respiratory system during embryonic development. As these organs are specified via a process of respiratory-esophageal separation (RES), the esophageal epithelium forms a simple columnar epithelium and then transforms into a stratified multi-layered epithelium [1–4]. The cellular and molecular mechanisms regulating RES and the esophageal epithelial morphogenesis during embryonic development have been extensively studied in recent years[2, 4–9]. However, it is less clear how the mature esophageal epithelium executes its self-renewal and maintenance of the proliferation-differentiation homeostasis.

Adult stem cells are vital for tissue/organ maintenance. Two models, the homogeneity and heterogeneity models, are proposed for self-renewal and maintenance of the proliferation-differentiation homeostasis of the mature esophageal epithelium[4, 10, 11]. The homogeneity model hypothesizes that cells in the basal layer consist of one single population that can function as the stem-like progenitors via the cell division cycle to produce daughter cells. Thus, these cells choose randomly to remain as progenitors or differentiate into suprabasal cells[12–15]. In contrast, the heterogeneity model offers an alternative possibility where, like other organs such as colon and stomach with a simple columnar epithelium or skin with stratified squamous epithelium, the basal layer of mammalian esophagi has a slow-cycling or quiescent stem cell subpopulation that can be self-renewal, giving rise to fast-dividing progenitor cells in the basal layer and/or all other differentiated lineages in the suprabasal layers and the differentiated layers[11]. In support of this model, asymmetrical cell division and cells with specific stemness related markers were found in the basal layers of mammalian esophagi[16–20]. Tissue-reconstitution and organoid formation indicated that cells isolated with various stemness related markers from mammalian esophagi could efficiently regenerate a completely stratified multi-layered squamous epithelium when compared with cells without these markers [21–24]. As single-cell RNA sequencing (scRNA-seq) pushed identification of cell populations at single cell resolution, Busslinger et al., recently identified a quiescent *Col17a1^high^ KRT15^high^* stem/progenitor cell population from the basal cell layer of human esophagi by scRNA-seq [25]. Hence, an accumulation of evidence supports the heterogeneity of the basal cells of mammalian esophagi. However, it remains unclear what proportion of basal cells are the stem cells, where the stem cells are located and how the stem cells are defined to maintain proliferation-differentiation homeostasis in the basal layer of the esophageal stratified squamous epithelium.

In this report, we show that 4-7% of slow cycling/quiescent basal cells (QBCs) with the molecular stem characteristics function as the stem cells spatially and randomly located in the basal layers of rat esophagi and normal rat esophageal keratinocyte cell line derived organoids. QBCs represent a unique cell population with unique patterns of DNA methylations and mRNA expressions. Detailed analyses indicate that high levels of hemidesmosomes (HDs) and low levels of Wnt signaling could sever as the critical determinants required for the stem cell maintenance and proliferation-differentiation homeostasis in rat esophageal stratified squamous epithelia.

## Results

### Identification of slow cycling/quiescent basal cells (QBCs) in mammalian esophagi

We sought to determine if a stem cell subpopulation could be detected in the basal layers of the mammalian esophagi. Esophageal tissues from rat, mouse and human were collected, fixed and stained with hematoxylin-eosin (H&E stain, S1 Fig A) and various cell markers (S1 Fig B-D). While H&E stain revealed the typical stratified squamous epithelia in rat, mouse and human esophageal tissues, the undifferentiated keratinocyte marker, cytokeratin14 (CK14), marked cells in the basal layer and the differentiated keratinocyte marker, cytokeratin13 (CK13) marked cells in the suprabasal layer and the differentiated layer. Stemness-related marker, P63, was detected in basal cells but not suparbasal cells in these tissues whereas other stemness-related markers, SOX2, BMI1 and OCT4 and DNA synthesis marker, PCNA, were stained in both basal and suprabasal cells. Previously, Neurotrophin receptor component P75NTR and hemidesmosome components integrin α6 (ITGα6) and β4 (ITGβ4) were indicated as potential stem cell markers of esophageal keratinocytes[18, 21]. Immunofluorescence staining showed that P75NTR, ITGα6 and ITGβ4 were clearly detected at cell membranes of basal cells (S1 Fig B and D). While P75NTR was stained ubiquitously in the basement membranes of basal cells, ITGα6 and ITGβ4 staining showed having subtle variations in the basement membranes of basal cells (S1 Fig B and D). However, quantifications of the staining intensities of ITGα6 and ITGβ4 were performed but the results were inconclusive. In sum, these results indicated that these makers could distinguish basal and/or suprabasal cells but could not precisely mark stem cells in mammalian esophagi.

**Figure 1.**
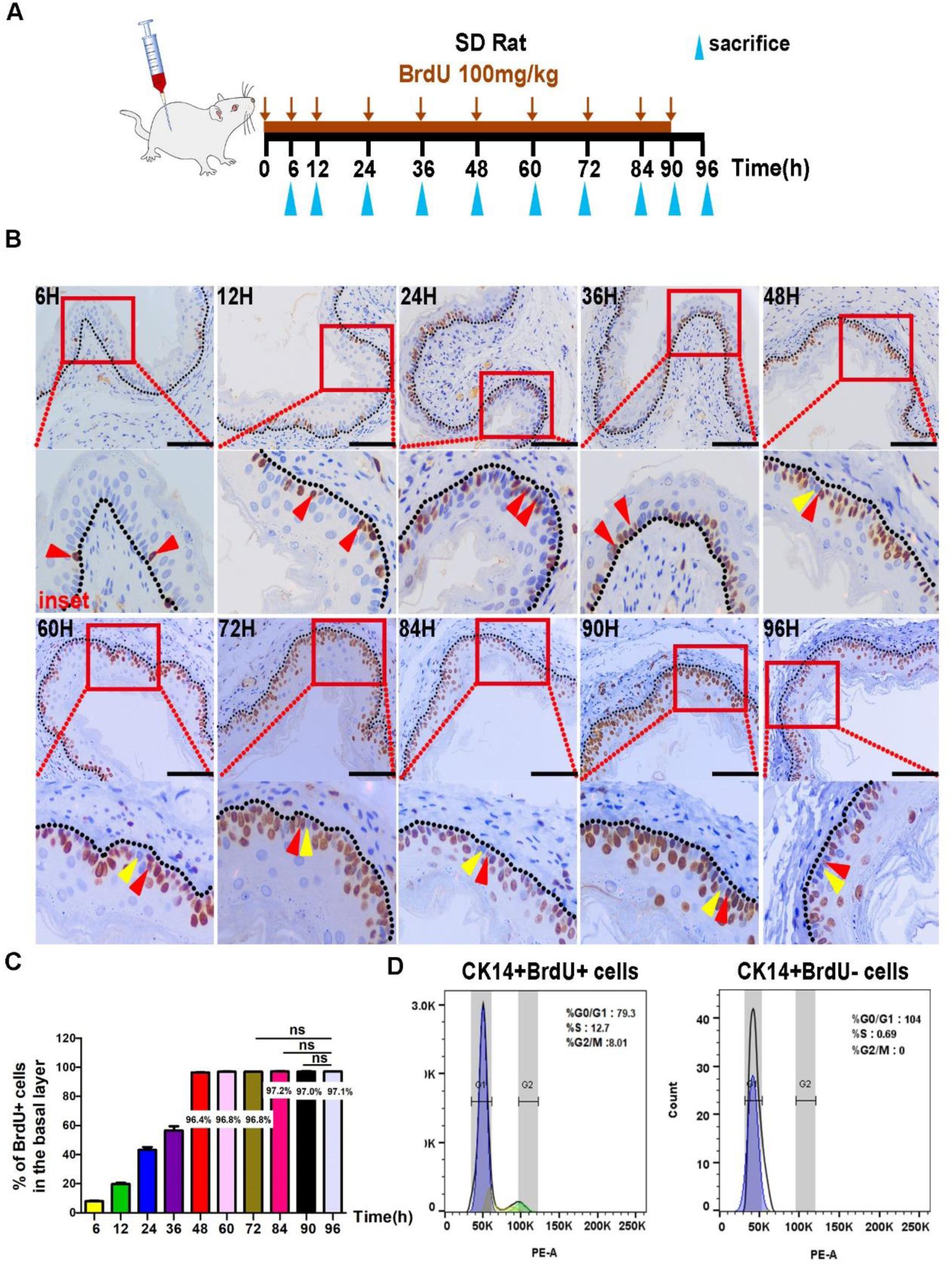
Rat esophageal basal layer exists a small relatively slow cycling/quiescent cell population. (A) Schematic illustration of BrdU-labeling experiment. SD rats were injected with BrdU of 100 mg/kg body weight once per 6 hours for 4 days and sacrificed at designed time points. (B) Immunohistochemistry staining of BrdU of the esophageal sections at designed time points. Red triangles indicate BrdU+ cells; the yellow triangles indicate BrdU-cells. Bottom panels represent the magnification of the interest regions indicated by a red rectangle of the top panels. (C) The percentage of BrdU+ cells in the basal layer of rat esophageal epithelium at designed time points. (n=5, n represents 5 intact basal layers of esophageal epithelium counted at each time point). (D) Cell cycle profile experiment of BrdU+ cells and BrdU-cells in the basal layer from 96 h label-chase rat esophagi by fluorescent-activating cell sorting (FACS). Dotted line marks the basement membrane. Scale bars: 100 μm. Data are represented as the mean +/-SD for percent analysis (*p < 0.05, **p < 0.01, ***p < 0.001)

Previous studies showed that stem cells in mammalian gastrointestinal track and skin were represented by a subpopulation of relatively slow-cycling/quiescent basal cells[26–29]. Long-term 5-Bromo-2’-deoxyuridine (BrdU) and 5-iodo-2’-deoxyuridine (IdU) label-retaining experiments indicated that slow-cycling/quiescent basal cells were also existed in the mouse and human esophageal epithelia[17, 18]. So, we took a shortcut and applied in vivo BrdU label-chase experiments to identify the potential stem cells in rat esophageal stratified squamous epithelia. A detailed experimental flowchart was shown in Figure 1A. Sprague-Dawley (SD) rats were given 100 mg/kg BrdU by intraperitoneal injection and BrdU labeled rat esophageal cross-sections were obtained at the indicated time points. These sections were stained with anti-BrdU antibody by immunohistochemistry or immunofluorescence analyses (Fig 1B and S2 Fig A-C). The label-chase experiments showed that BrdU labeled cells (BrdU+) were identified mainly in the basal layer in short-labeling times and then detected in the basal layer and the suprabasal layers in long-labeling times (>48 h). Cell counting indicated that BrdU+ cells in the basal layer increased gradually and, ultimately, reached to a maximum level at 48 h labeling time point (Fig 1B and S2 Fig C). Labeling BrdU up to 72-96 h, BrdU+ cells remained at the same level in the basal layer (Fig 1C and S2 Fig C). The BrdU+ and BrdU-(BrdU unlabled) cells were ∼96% and ∼4%, respectively. Similar results were also obtained from BrdU label-chase experiments in BALB/c mice (S2 Fig D and F).

**Figure 2.**
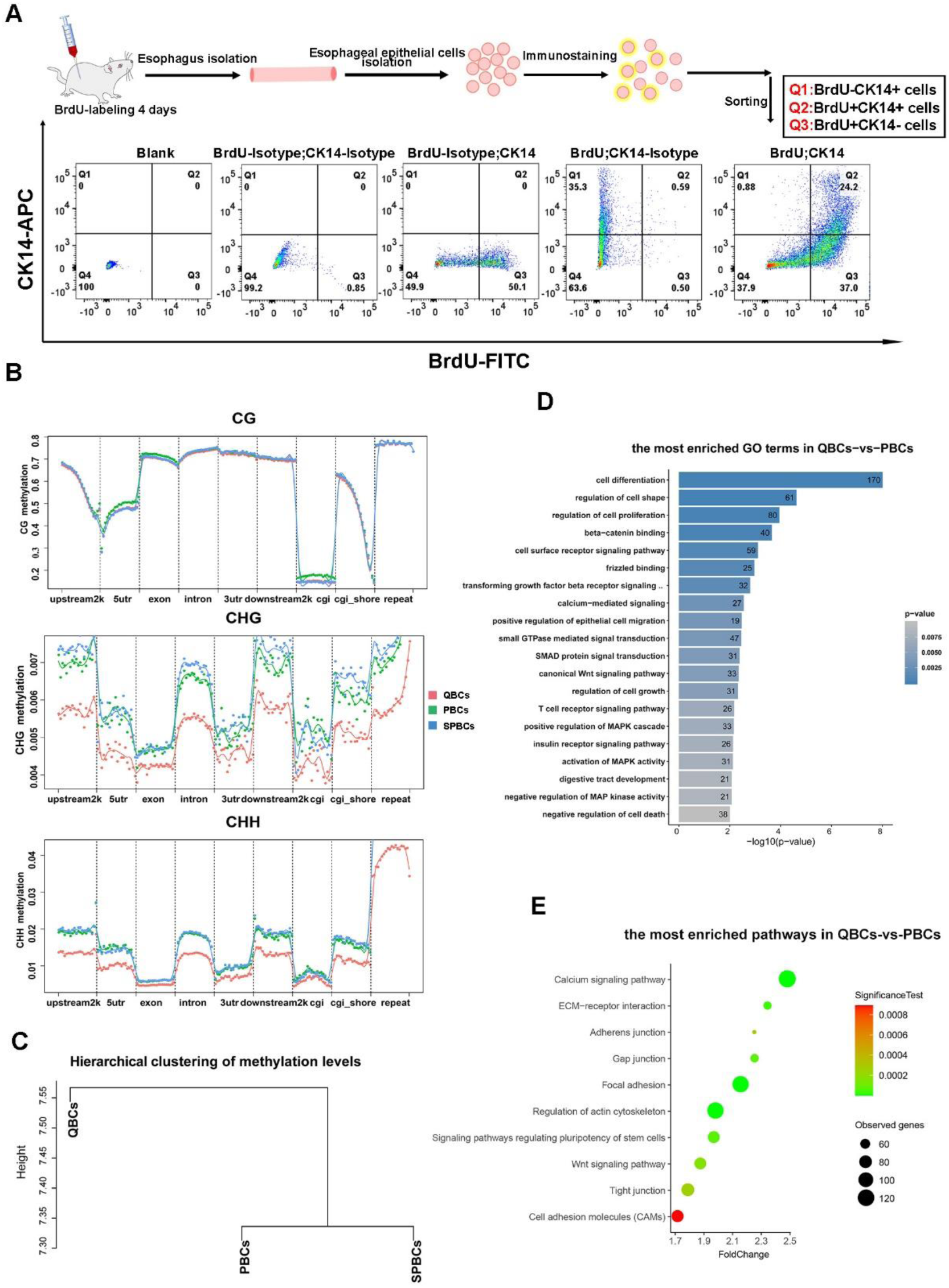
Quiescent basal cells (QBCs) population in the rat esophageal basal layer have a distinct methylation profile. (A) Three populations, BrdU-CK14+/quiescent basal cells (QBCs), BrdU+CK14+/proliferating basal cells (PBCs) and BrdU+CK14-/suprabasal proliferating cells (SPBCs), were sorted from the esophageal keratinocytes of SD Rats with BrdU-labeling for 96 hours. (B) Clustering analysis of CpG, CpHpG and CpHpH (H=A, C and T) methylation levels among three populations. (C) Hierarchical clustering in the methylation maps among three populations. (D) Histogram showing the most enriched GO term in QBCs-vs-PBCs. (E) Bubble plot showing the most enriched KEGG pathways in QBCs-vs-PBCs.

We determined the cell cycle profiles of BrdU+ and BrdU- cells isolated from 96 h label-chase rat esophagi by fluorescent-activating cell sorting (FACS). The results demonstrated that, indeed, BrdU+ cells were cycling cells whereas BrdU- cells were slow-cycling/quiescent cells at the G0/G1 phase of the cell cycle (Fig 1D). Double staining of BrdU with potential esophageal stemness markers, cytokeratin 15 (CK15), ITGα6, CD34, or P75NTR, and stemness-related markers, SOX2, BMI1 or OCT4, revealed that, consistent with the results shown in Figure S3, the individual cell markers did not specifically and sufficiently mark BrdU+ cycling cells or BrdU- slow cycling/quiescent cells in the basal layer (S3 Fig B and C). However, we noticed that all BrdU- slow cycling/quiescent cells in the basal layer were co-expressed the stemness-related markers SOX2, BMI1 and OCT4 (S3 Fig D and E). We determined the distribution patterns of BrdU- cells but found that they were randomly located in the basal layer. Taken together, these results indicated that the basal layer of the mammalian esophagus contained slow cycling/quiescent basal cells (QBCs). However, these cells could not be specifically and effectively marked by current known stemness related markers.

**Figure 3.**
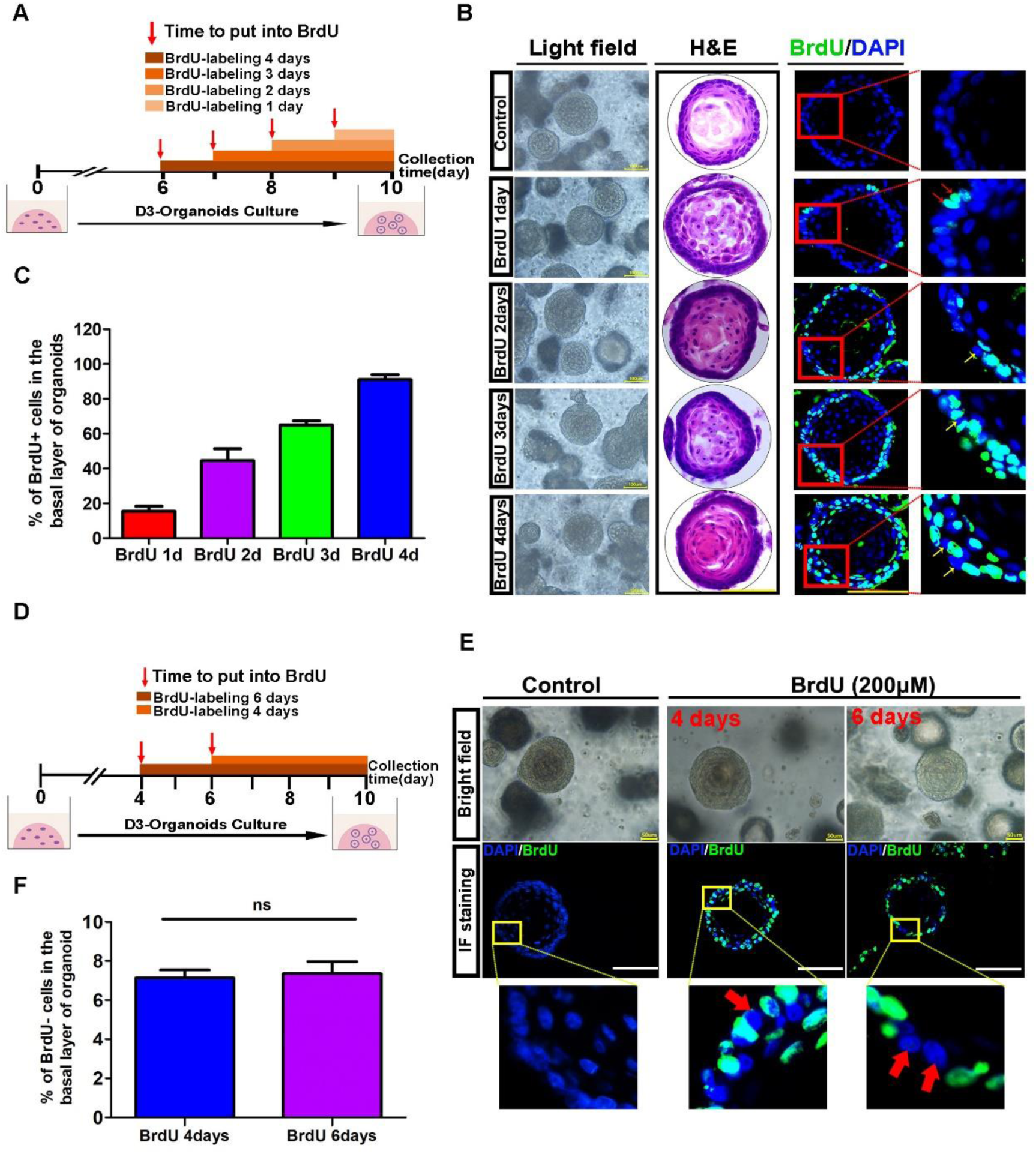
The basal layer of rat esophageal organoids derived from D3 cells also exists quiescent basal cells (QBCs) population. (A) Schematic illustration of BrdU-labeling experiment of rat esophageal organoids derived from immortalized normal esophageal keratinocyte cell line D3. 200μM of BrdU concentration was used at different time points in the organoid culture process. (B) The representative images of light field, HE staining and corresponding BrdU immunofluorescence staining of D3-organiods culturing at different BrdU labeling days. (C) The percentage of BrdU+ cells in the basal layer of D3- organoids culturing in different BrdU labeling days. (n=9, n presents nine random microscope fields, 400x). (D) Schematic illustration of BrdU-labeling at 4 days and 6 days of D3-organoid culture. (E) The representative images of light field and immunofluorescence staining of BrdU of D3-organoids for BrdU- labeling at 4 and 6 days. Red arrows indicate BrdU- cells in the basal layer of D3-organoids. (F) The percentage of BrdU- cells in the basal layer of D3- organoids at BrdU labeling 4 and 6 days. (n=9, n presents nine random microscope fields, 400x). Scale bars: 100μm. Data are represented as the mean +/- SD for percent analysis (*p <0.05, **p < 0.01, ***p < 0.001).

### Determination of QBCs as a unique cell population by genome-wide DNA methylation profiling

To further define molecular characteristics of QBCs, we decided to perform whole genome bisulfite sequencing (WGBS) profiling experiments as transcriptome analyses could not be done with cells fixed and stained by BrdU antibody. Rats were labeled with BrdU for 4 days and then esophageal epithelial cells were sorted into three populations, BrdU-/CK14+, BrdU+/CK14+ and BrdU+/CK14-, based on BrdU and CK14 antibody staining by FACS (Fig 2A and S3 Fig A). Consistent with immunostaining results (Fig 1C), BrdU-/CK14+ cells represented about 4% of CK14+ cells, which were consistent with the immunohistochemistry or immunofluorescence analyses. We renamed BrdU-/CK14+ cells as QBCs (quiescent basal cells), BrdU+/CK14+ cells as PBCs (proliferating basal cells) and BrdU+/CK14- cells as SPBCs (suprabasal cells). WGBS profiling showed that overall methylation levels of QBCs, PBCs and SPBCs of Rattus norvegicus were similar, especially at CpG sties, which covered >70% methylation sites detected (S4 Fig A-C). However, detailed analyses of the cytosine methylations at none CpG sites, such as CpHpG sites and CpHpH sites (H=A, C and T), revealed different results in QBCs, PBCs and SPBCs. The none CpG methylations in QBCs were significantly lower than those in PBCs and SPBCs (Fig 2B). More importantly, although overall methylation levels of QBCs, PBCs and SPBCs were similar, hierarchical clustering in the methylation sites based on genomic sequence could clearly separate QBCs from PBCs and SPBCs, indicating that epigenetic regulations at DNA levels were distinct in QBCs when compared with those in PBCs and SPBCs (Fig 2C).

**Figure 4.**
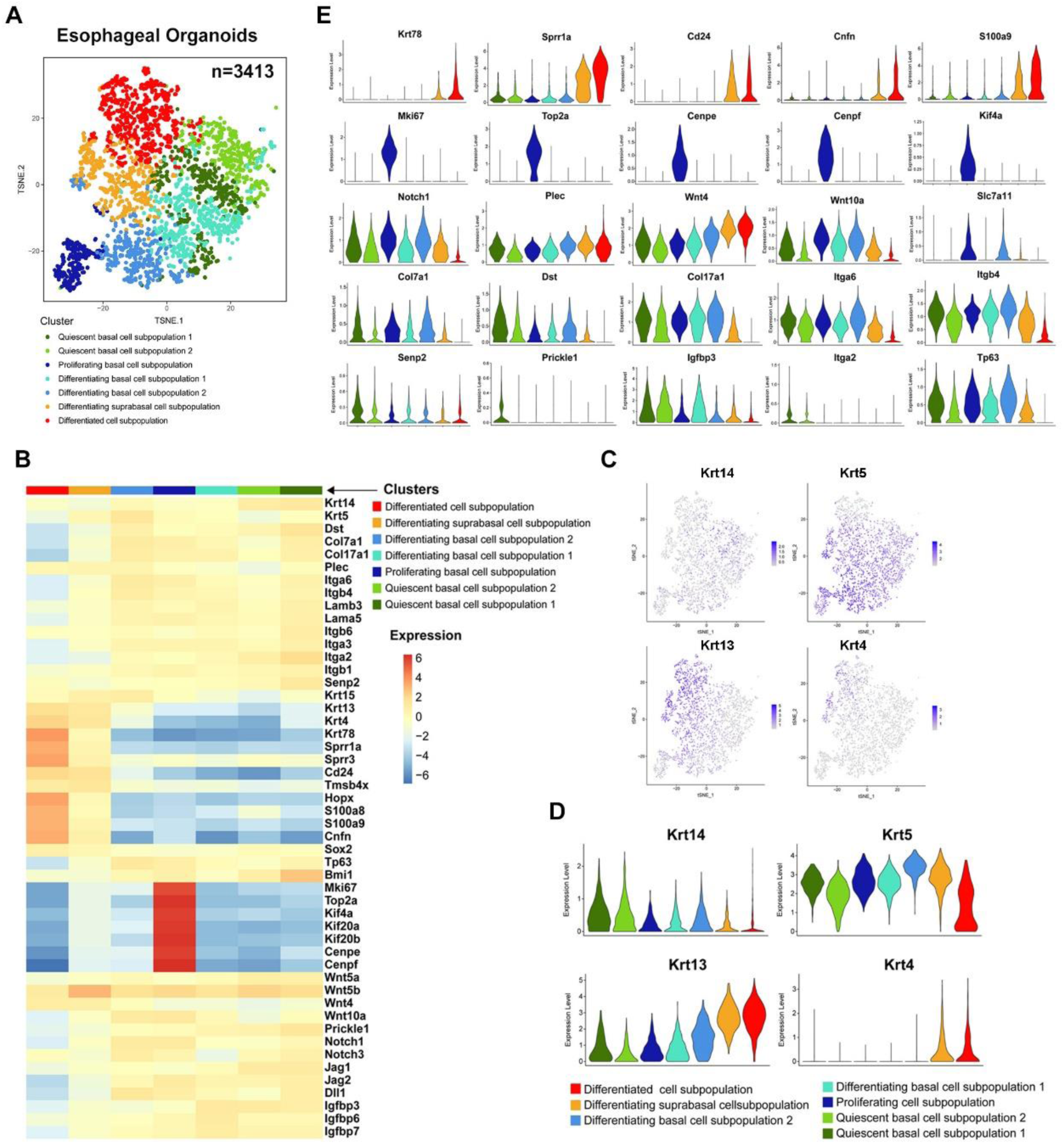
Single cell RNA sequencing analysis of rat esophageal organoids derived from D3 cells. (A) t-SNE plot displaying the scRNA-seq data of rat D3-organoids. Different colors indicated the distinct cell subpopulations. (B) Heatmap showing the selected genes expression from the 7 clusters corresponding to the subpopulations of D3-organoids. (C) UMAP plots of Keratin genes expression among the 7 different subpopulations of D3- organoids. (D) Violin plots of Keratin genes expression among the 7 different subpopulations of D3-organoids. The y-axis represents the expression level of genes, and the x-axis represents different subpopulations. (E) Violin plots of selected genes expression levels among 7 different subpopulations.

We defined differentially methylated regions (DMRs) between QBCs and PBCs. Consistent with the results obtained from hierarchical clustering (Fig 2C), heatmaps generated from DMR methylation levels in top-scored gene surround regions clearly displayed the differences in QBCs and PBCs (S4 Fig D and E). Gene Ontology (GO) and Kyoto Encyclopedia of Genes and Genomes (KEGG) enrichment analyses were performed in genes with DMRs. As shown in Fig 2D, the highly enriched GO terms were mainly associated with biological processes, such as cell differentiation, regulation of cell shape and regulation of cell proliferation, etc. The highly enriched KEGG terms were mainly associated with genes involved in regulating focal adhesion, cytoskeletal structure and regulation, adhesion junction and Wnt signaling pathway (Fig 2E). These results indicated that even though QBCs and PBCs were spatially closed in the basal layer of the esophagus, their DNA methylation levels and patterns in genome, especially in gene surrounding regions that controlled gene expressions, were different. The epigenetic characteristics at DNA level (DNA methylations) in QBCs was unique when compared with those of PBCs and SPBCs, thus representing a cell population located spatiotemporally in the esophageal basal layer.

### Existence of the QBCs in normal rat esophageal keratinocyte cell line derived organoids

Three-dimension (3D) organoid cultures have been demonstrated as a powerful in vitro system for studying identification, determination, self-renewal trajectory and maintenance of tissue stem cells, formation and proliferation-differentiation homeostasis of organ tissues and carcinogenesis [9, 30–32]. To further characterize QBCs, we generated a human telomere reverse transcriptase (h-tert) immortalized rat normal esophageal keratinocyte cell line (RNE-D3, for detail see Experimental procedures). Like rat esophageal primary epithelial keratinocytes, REN-D3 (D3) cells in Matrigel with conditional culture medium formed normal and typical esophageal organoids with endodermal morphological structures including the basal layer, the suprabasal layers and the keratin layer in 10 days (S5 Fig A-C).

**Figure 5.**
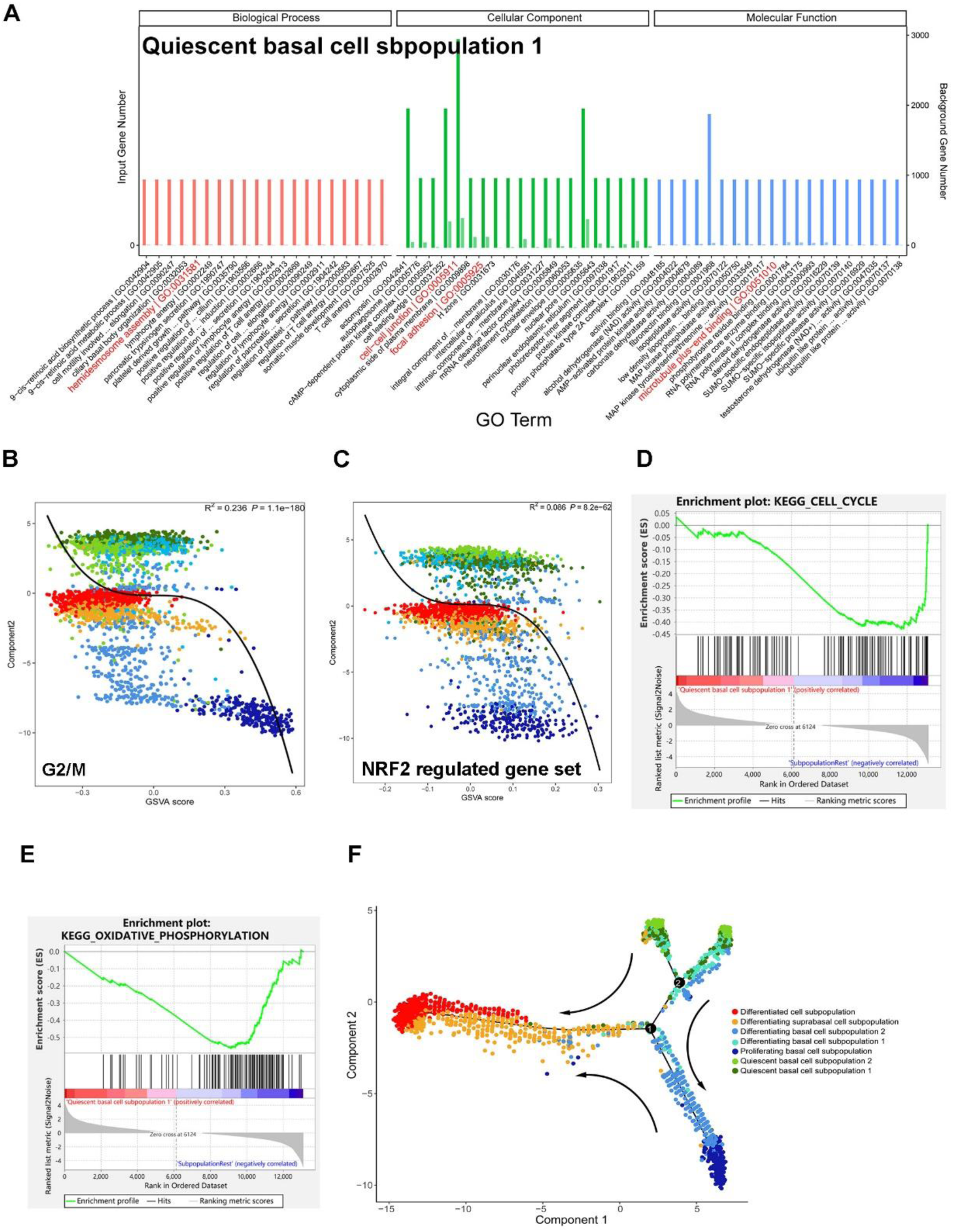
Detailed signatures of QBC1 of single cell RNA sequencing analysis. (A) GO enrichment analysis in QBC1 (quiescent basal cell subpopulation 1) showed significant upregulation of hemidesmosome component, in accordance with related cell adhesion and cytoskeleton change. (B) GSVA of genes that controlled the cell cycle progression in 7 different subpopulations of D3-organoids. (C) GSVA of Nrf2 regulated gene set in 7 different subpopulations of D3-organoids. (D) And (E) GSEA showed significant downregulation of cell cycle regulating genes (upper panel) and oxidative phosphorylation regulating genes (lower panel) in QBC1 (quiescent basal cell subpopulation 1). (F) Pseudotime trajectory ordered 7 different subpopulations of D3-organoids in a two-dimensional state-space. The x and y axes are two principal components. The numbers in the black circles represent nodes that determine different cell states in the trajectory analysis. The black arrows indicate the evolutions of cell fates.

Similar to rat esophageal stratified squamous epithelia, immunofluorescence assay showed that CK14, P75NTR, ITGα6 and ITGβ4 were expressed in the basal layer cells while PCNA, BMI1, SOX2 and OCT4 were expressed in the basal and suprabasal layer cells and CK13 was expressed in the suprabasal layer cells and the keratin layer cells in D3 derived organoids (D3-organoids). These results indicated that D3-organoids could be used as a convenient in vitro model for esophageal study (S5 Fig D and E).

BrdU label-chase experiments were applied to determine if QBCs were presented in D3-organoids. The concentration of BrdU used in the experiments were determined as 200 μM, which could sufficiently and effectively label cells but not inhibit DNA synthesis in D3-organoids (S6 Fig A-D). Detailed experimental flowchart was shown in Figure 3A. Growing D3-organoids from day 6 to day 9 were labeled with BrdU, respectively, and then the labeled organoids were collected, fixed and analyzed at day 10. Immunostaining showed that D3-organoids labelled with BrdU for 1, 2, 3 and 4 days displayed 16%, 42%, 69% and 93% of BrdU+ cells in the basal layer, respectively (S3 Fig B and C). To ensure BrdU labeled cells in the basal layers of D3-organoids reached maximum, we prolonged BrdU labeling time in growing D3-organoids, starting at day 4 for 6 days (Fig 3D). The results showed that, like the results obtained from D3-organoids labeled with BrdU for 4 days, BrdU labeling in growing D3-organoids for 6 days did not further increase BrdU+ cells in their basal layers. BrdU+ cells were about 93% and BrdU- cells were about 7% in the basal layer (Fig 3E and F). Hence, BrdU label-chase experiments demonstrated that, akin to rat esophageal stratified squamous epithelia, D3- derived normal esophageal organoids also contained QBCs in the basal layer.

**Figure 6.**
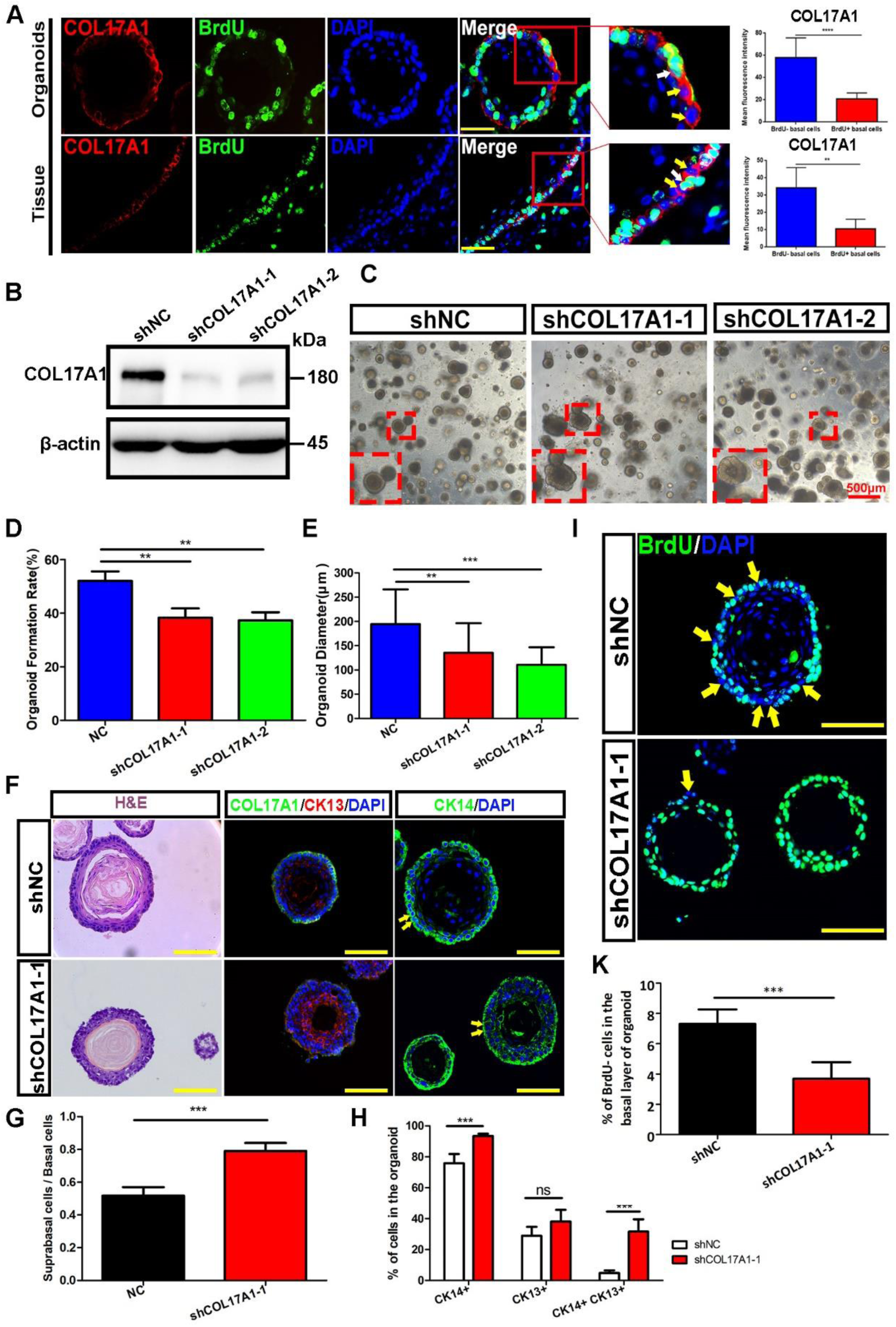
Hemidesmosome (HD) component, COL17A1, in stem cell maintenance and proliferation-differentiation homeostasis of rat esophagus and organoids. (A) COL17A1 expression were significantly higher in BrdU- basal cells than that in BrdU+ basal cells in D3 organoids and esophageal tissue. White arrow and yellow arrow stand for representative BrdU+ and BrdU- basal cells with COL17A1 expression discrepancies, respectively. Histograms on the right panel displayed quantifications of COL17A1 florescence intensity using Fiji ImageJ. Scale bars: 50μm. (B) Western Blotting verification of D3-sh*Col17a1* cell line construction. (C) The representative images of light field of organoids at day 10. COL17A1 knockdown significantly inhibited organoid formation and growth. Scale bars: 500μm. (D) Quantification of organoid formation rate after COL17A1 knockdown. (E) Quantification of organoid diameter after COL17A1 knockdown. (F) H&E staining and immunofluorescence staining of intermediate filaments (CK13 and CK14) of organoids showed significant self-organization perturbation presented as uneven basal layers and abnormal distribution of CKs after COL17A1 knockdown. (G) The ratio of suprabasal cells vs. basal cells of D3-organoids after COL17A1 knockdown. (n=5, n presents six random microscope fields,400X). (H) The percentage of CK14+ cells、CK13+ cells and CK14+CK13+ cells of D3-organoids after COL17A1 knockdown. (n=5, n presents six random microscope fields,400X). I) Immunofluorescence staining of BrdU of D3-organoids labeled for 4 days after COL17A1 knockdown. The yellow arrows indicated the BrdU- cells. Scale bars: 100μm. (J) The percentage of BrdU- cells in the basal layer of D3-organoids after COL17A1 knockdown. (n=6, n presents six random microscope fields,200X). Data are represented as the mean +/- SD for percent analysis (*p < 0.05, **p < 0.01, ***p< 0.001).

### Resolution of QBCs at the single-cell level in rat esophageal organoids

We performed single-cell RNA sequencing (scRNA-seq) in D3-organoids to identify QBCs with the distinct RNA expression patterns. D3-organoids grown for 10 days were collected, digested into single-cell suspension, and then processed for scRNA-seq using the sorting and robot-assisted transcriptome sequencing (SORT-seq) protocol. The scRNA-seq analysis resulted in identification of 3413 keratinocytes from D3-organoids that could be classified into 7 distinct but related subpopulations (Fig 4A). Based on the expression of known keratinocyte undifferentiation-differentiation markers, *CK14* (also called *Krt14*) and *CK13* (also called *Krt13*), D3-organoid epithelial cells could be first categorized as two cell populations, *CK14^high^* population that represented basal cells and *CK13^high^* population that represented suprabasal cells and differentiated cells (Fig 4B-D). *CK13^high^* population could be further clustered into suprabasal cell subpopulation with high levels of *Sox2*, *Bmi1* and *CK5* (also called *Krt5*) gene expression and differentiated cell subpopulation with high levels of *Krt78*, *Sprr1a*, *Cd24*, and *Cnfn* differentiation-related gene expression (Fig 4B and 4E). Moreover, *CK14^high^* population could be clustered into 5 subpopulations where one subpopulation expressed high levels of the cell cycle related genes, such as *Ki67*, *Top2A*, *Kif4A*, *Cenpe* and *Cenpf* (Fig 4B and 4E). Although the other four subpopulations all expressed low levels of the cell cycle related genes, two of the subpopulations also expressed high levels of differentiation-related genes, *Notch1*, and Wnt signaling components, *Wnt4 and Wnt10a* (Fig 4B and 4E). These two subpopulations could be further separated based on expression levels of proliferation related gene, *Igfbp3* (Fig 4B and 4E), and ROS (reactive oxygen species) related gene, *Slc7a1* (Fig 4E). In contrast, the rest of two subpopulations expressed high levels of Wnt signaling negative regulators, *Senp2* and *Prickle1* and basement membrane markers, *Col17a1* (also called *Bpag2*), *Dst* (also called *Bpag1*), *Itgα6*, *Itgβ4* and *Col7a1* genes (Fig 4B and 4E). However, one subpopulation expressed these basement membrane markers higher than the other.

Based on the results, we named 7 cell subpopulations as QBC1 (quiescent basal cell subpopulation 1), QBC2 (quiescent basal cell subpopulation 2), PBC (proliferating basal cell subpopulation), DBC1 (differentiating basal cell subpopulation 1), DBC2 (differentiating basal cell subpopulation 2), DSC (differentiating suprabasal cell subpopulation) and DC (differentiated cell subpopulation) (Fig 4). We focused on QBC1/2 for further analyses where GO and GSVA (gene set variation analysis) for multi-biological functions were determined these subpopulations [33, 34]. The results showed that the significantly enriched GO terms in QBC1/2, especially QBC1, were cell differentiation, regulation of cell shape and regulation of cytoskeletal structure, such as hemidesmosome (HD) assembly, cell-cell junction, focal adhesion and microtubule plus-end binding (Fig 5A).

GSVA demonstrated that QBC1, QBC2 as well, displayed significant low expressions in genes that controlled the cell cycle progression, ROS and their related phosphorylation regulation pathways when compared with other cell subpopulations, especially PBC (Fig 5B and 5C). GSEA (gene set enrichment analysis) also indicated that QBC1, significant down-regulated of cell cycle and oxidative phosphorylation related genes (Fig 5D and 5E). These results consistent with the cell cycle profiles of QBCs which isolated from rat esophagi by FACS that we mentioned in Figure 1D. The results also demonstrated that QBC1/2 with low cell cycle genes expression pattern led to the phenotype of slow-cycling/quiescent cells at the G0/G1 phase of cell cycle, suggesting that QBC1/2 identified in scRNA-seq and QBCs identified by BrdU label-chase experiments represented the same stemness population in the basal layer of esophagus.

Identification of 7 subpopulations from D3-organoids allowed us to perform pseudo-time cell trajectory and determine evolutions of esophageal epithelial cell fates during proliferation-differentiation homeostasis. As shown in Figure 5F, cells in QBC1/2 cells could be positioned as the start points. Pseudo-time cell trajectory indicated that, as the most stemness subpopulation in esophageal basal layer, QBC1/2 produced DBC1 and DBC2, which progressed into DSC, ultimately, differentiating into DC or QBC1/2 generated PBC cells that, in turn, progressed into DSC and finally differentiated into DC cells.

In sum, at the single-cell level, we identified 7 subpopulations in rat esophageal organoids, with individual mRNA expression patterns, 2 subpopulations were in suprabasal layers whereas 5 subpopulations, including the most stemness subpopulation QBC1/2, were in basal layer, that indicating the heterogeneity of mammalian stratified squamous epithelium, especially the basal layer of esophagus.

### High levels of HD components marked QBCs in rat esophageal epithelia and organoids

Since QBC1/2 and QBCs represented the same stemness population in the basal layer of esophagus, we compared the results obtained from scRNA-seq and WGBS, especially genes identified by scRNA-seq in QBC1/2 and WGBS in QBCs. The results from two sequencing data showed that QBC1 was enriched the expressions of HD components (*Itgβ4*, *Col17a1*, *Dst* and *PLEC*), HD-anchoring extracellular matrix *Lamb3*, and Wnt pathway negative regulator, *Prickle1* (Fig 4B), while hypomethylated sites were also detected in the sequences of these genes and/or the surrounding regions in QBCs (S4 Fig E). Consistent with these results, recent studies showed that the maintenance of Wnt pathway at low level could facilitate the specification of anterior foregut endoderm toward the esophageal progenitor cells lineage[9, 31] while COL17A1 was required for skin keratinocyte stem cell maintenance[35]. High expression of COL17A1 was also identified as a marker of human esophageal quiescent stem/progenitor cells[5]. Hence, these results suggested that high levels of HD components and Wnt pathway negative regulators could sever as prominent markers of QBCs in the basal layer of the esophageal epithelium.

To define if HD and Wnt signaling components marked QBCs in the basal layers of the rat esophagi, we examined their subcellular localizations in the basal layers of the rat esophageal epithelia and D3-organoids. While HD components, ITGα6 and ITGβ4, staining showed strong fluorescence in the basement membranes of basal cells (S1 Fig B and D), another HD component PLEC and Wnt pathway negative regulator Prickle1 were stained in both basal and suprabasal cells (S1 Fig E). In contrast, other HD component, COL17A1, displayed obviously cell staining variations in the basement membranes of basal cells (S1 Fig E).

To determine if COL17A1 marked QBCs, we examined the expression and subcellular localization of COL17A1 in the basal layers of the esophagi of rats labeled with BrdU for 4 days. High levels of COL17A1 were detected in QBCs when compared with those in PBCs of esophageal epithelia tissues (Fig 6A). Similar results were also obtained in esophageal epithelia D3-organoids labeled with BrdU for 4 days (Fig 6A). Notably, high levels of DST, another component of HD, were also detected in QBCs when compared with those in PBCs (S8 Fig A and B). Thus, these results demonstrated that HD components, COL17A1^high^ and DST ^high^ could be used as prominent markers to mark stem cells in the basal layer of the esophageal epithelium.

### Involvements of hemidesmosome (HD) and/or Wnt signaling in stem cell maintenance and proliferation-differentiation homeostasis in rat esophageal organoids

To ascertain that high levels of HD components were not only prominent markers for stem cells but also involved in regulating stem cell maintenance and proliferation-differentiation homeostasis in the basal layer of the esophagus, we perturbed HDs via RNAi depletion in D3 cells and examined organoid formation in HD depleted D3 cells. We first knocked down expression of COL17A1 in D3 cells by transduction of lentivirus expressing *Col17a1* shRNA. Immunoblotting analysis showed that lentivirus expressing *Col17a1* shRNA in D3 cells could effectively and efficiently ablate endogenous COL17A1 expression (Fig 6B). As shown in Figures 6C-E, when COL17A1 depleted D3 cells and controls were grown in Matrigel with conditional culture medium for 10-12 days and analyzed for organoid formations, ablation of COL17A1 expression in D3 cells significantly affected organoid formations including inhibition of organoid formation rate, reduction of organoid size, perturbation of organoid shape and destruction of proliferation-differentiation homeostasis. Immunofluorescence with anti-CK14 and anti-CK13 antibodies or immunohistochemistry with H&E stain showed that, when compared with control D3-organoids, COL17A1 depleted D3-organoids grew and developed aberrantly, forming disorganized stratified epithelia with non-smooth budding sharps, reduced sizes and abnormal CK14 and CK13 staining (Fig 6F). The ratio of suprabasal layer cells vs basal layer cells was increased significantly in COL17A1 depleted D3-organoids when compared with controls, indicating an imbalance in proliferation-differentiation homeostasis during organoid formation and growth (Fig 6G). In addition, CK14 positive cells was also increased in COL17A1 depleted D3-organoids, suggesting that the imbalance homeostasis might be caused by the insufficient differentiation (Fig 6H). Therefore, we performed BrdU labeled experiments to measure the QBCs (BrdU- cells) in the basal layers of COL17A1 depleted D3-organoids. The results showed that, when compared with 7% of QBCs in D3-organoids labeled with BrdU for 4 days, the QBCs in COL17A1 depleted D3-organoids were significantly reduced to ∼4% in the basal layer (Fig 6I and 6J). These results demonstrated that COL17A1, a core component of HD and a QBCs marker, was required for QBC maintenance and esophageal keratinocyte organoid formation.

Next, we ablated another HD component, PLEC, which was not only required for type I but also required for type II of HD formation, in D3 cells by RNAi and examined D3-organoid formation[36]. Consistent with results obtained from COL17A1 depletion, depletion of PLEC resulted in inhibition of the D3-organoid formation rate, reduction of the organoid size, perturbation of the organoid shape and disruption of proliferation-differentiation homeostasis (S7 Fig C-G). The QBCs in PLEC depleted D3-organoids were significantly reduced to ∼3% in the basal layer (S7 Fig H and I).

We next examined the role of Wnt signaling in regulating QBCs, the stem cell maintenance and proliferation-differentiation homeostasis in the basal layer of D3-organoids. Since Wnt activities in organoids were difficulty to measure directly, we performed functional assays to determine whether perturbations of Wnt activities would affect stem cell maintenance and proliferation-differentiation homeostasis in the basal cells. To this end, we perturbed Wnt activities by additions of Wnt pathway inhibitors IWP-2[37, 38] and SFRP-2[39, 40] or activator CHIR99201[41–43] in organoid culture media and examined organoid formation and morphogenesis. Detailed experimental flowchart was shown in Figure 7A. Growing D3-organoids at day 6 were treated with BrdU plus DMSO (control) or BrdU plus Wnt inhibitor, IWP-2 (2uM) or SFRP-2 (5nM), for additional 4 days and then the organoid morphogenesis and the proportion of QBCs in the basal layer cells were determined by histochemistry analysis or immunofluorescence using anti-BrdU antibody. H&E staining showed that although organoids in control or in Wnt inhibitor treatment obtained at day 10 could differentiate into stratified squamous tissue structures with the basal layer, the suprabasal layer and the differentiated layer, organoids treated with Wnt inhibitors were smaller in size than controls (Fig 7B and C). The BrdU label-chase experiments indicated that, when compared with control, D3 organoids treated with Wnt inhibitors, IWP-2 and SFRP-2, displayed dramatically increased levels of QBCs cells in the basal layers (∼7% in controls vs ∼16-18% in treatment with IWP-2 or SFRP-2) (Fig 7B and D). Consistently, quantitative RT-PCR (qRT-PCR) analysis of mRNA expressions of Wnt pathway target genes *Axin2* and *Dvl1* in the basal cells of D3-organoids treated with Wnt inhibitors or controls, which were isolated by FACS with anti-ITGβ4 antibody, demonstrated that *Axin2* and *Dvl1* expressions were significantly inhibited by Wnt inhibitor treatments (S7 Fig 7A-C). In contrast, treatment of D3-organoids with Wnt pathway activator CHIR99201 induced morphologic pre-mature differentiations of organoids when compared with control determined by H&E staining (S8 Fig C). Taken together, these results indicated that low levels of Wnt signaling benefited the maintenance of QBCs in the basal layer, required for the stem cell maintenance and proliferation-differentiation homeostasis in the esophageal stratified squamous epithelium.

**Figure 7.**
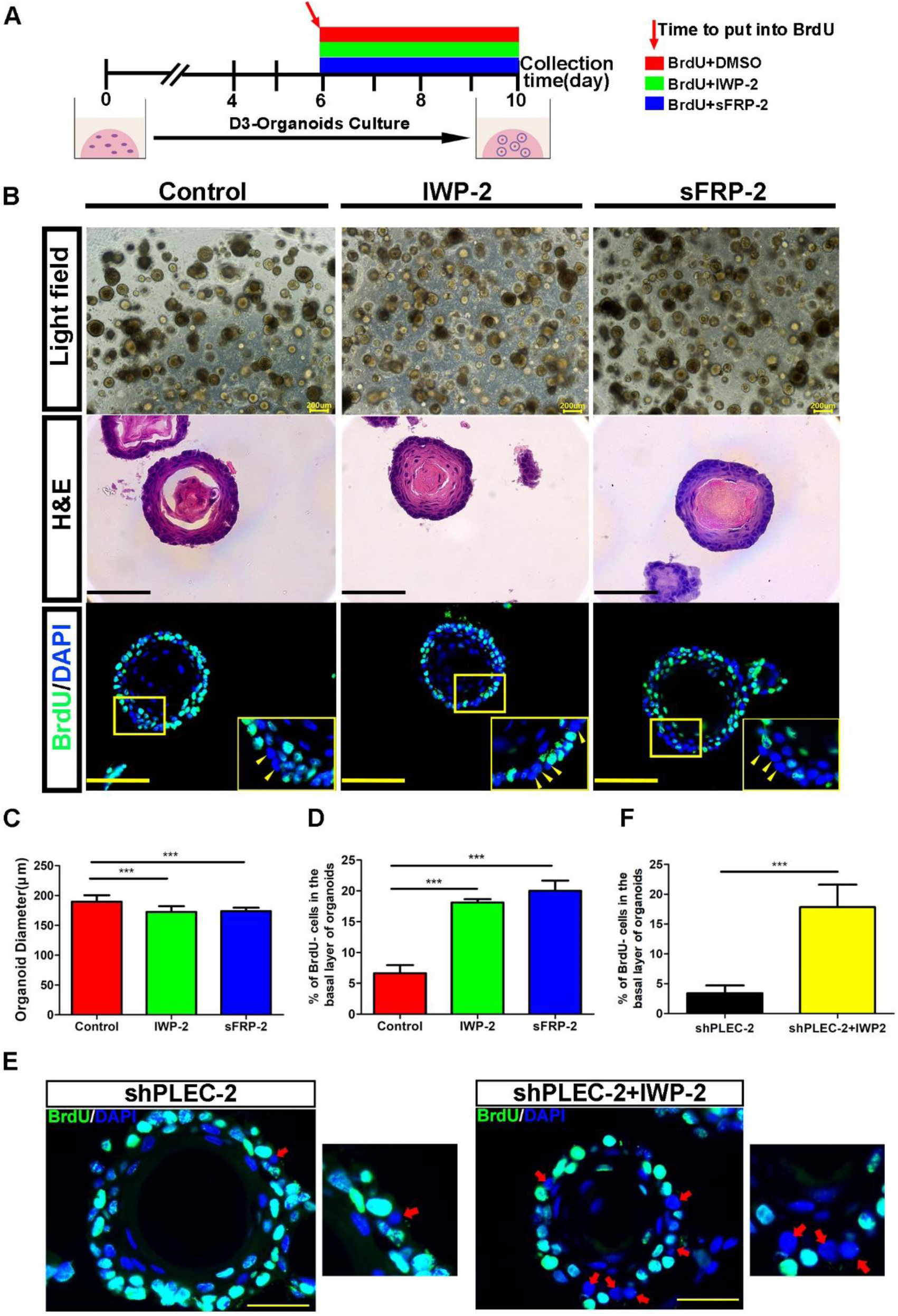
Wnt signaling-HD crosstalk can maintain the stem cell identity of quiescent basal cells in esophageal organoids. (A) Wnt inhibitors IWP-2 and sFRP-2 were added on the 6th day of esophageal organoid culture and BrdU-labeling experiment was performed at the same time. 200μmol BrdU, 2μmol IWP-2 and 5nmol sFRP-2 were used. (B) The images of light field, HE staining and corresponding BrdU immunofluorescence staining in (A). (C) The diameters of esophageal organoids in the groups with Wnt inhibitors IWP-2 and sFRP-2. (n=5, n presents five random microscope fields,200x). (D) The percentage of BrdU- cells in the outmost basal layer of esophageal organoids in the groups with Wnt inhibitors IWP-2 and sFRP-2. (n=10, n presents ten random microscope fields,400x). (E) Immunofluorescence staining of BrdU of D3-organoids after PLEC knockdown or/and treated with Wnt inhibitor IWP2 for 4 days. Scale bars: 100μm. (F) The percentage of BrdU- cells in the basal layer of D3-organoids after PLEC knockdown or/and treated with Wnt inhibitor IWP-2 for 4 days. (n=6, n presents six random microscope fields,400X). Data are represented as the mean +/- SD for percent analysis (*p < 0.05, **p < 0.01, ***p< 0.001).

Finally, we explored whether there was a possible cross-talk and/or interplay between HDs and Wnt signaling in controlling the stem cell maintenance and proliferation-differentiation homeostasis in the esophageal stratified squamous epithelium. Therefore, we ablated PLEC expression in D3-organoids and simultaneously treated the organoids with Wnt inhibitor, IWP2. When compared with D3-organoids depleted PLEC alone, IWP2 treatment significantly rescued the amount of QBCs in the basal layer of D3-organoids (Fig 7E and 7F). These results indicated that not only high levels of HDs and low levels of Wnt signaling but also a cross-talk and/or interplay between HD and Wnt signaling defined QBCs of the basal layer, which were crucial for the stem cell maintenance and proliferation-differentiation homeostasis in the mammalian esophageal epithelium.

## Discussion

Despite of extensive research, identification and determination of stem cells in the mammalian esophageal epithelia remained controversial[11]. By genetic means in mouse model system, Doupe et al., showed that the esophageal epithelium contained a single population of cells that divided stochastically to generate proliferating and differentiating daughter cells with equal possibility, thus indicating that a “reserve” slow cycling stem cell pool was not existed in the esophageal epithelium [13]. However, analyses of the human and mouse esophageal epithelium by histological and label-retaining analyses demonstrated existences of asymmetrically dividing, slow cycling stem cells in the basal layers of the esophageal epithelia[20, 24]. Furthermore, high expressions of many stemness markers, such as SOX2, ITGα6 and ITGβ4, were postulated to present in the stem cells of mammalian esophageal stratified squamous epithelia[21, 23]. Although several studies suggested that high expressions of these potential stemness markers in esophageal keratinocytes promoted cell stemness, the results were often inconclusive and controversial, thus meriting further investigations[4, 11].

Promoted by the label-retaining experiments where, in the absence of esophageal stem cell markers, long term (>3 months) tracking DNA syntheses/cell divisions by BrdU/IdU label-retaining in rodent and human esophageal stratified squamous epithelial identified slow-cycling/quiescent stem cell population[19, 24], we decided to take a shortcut by performing label-chase experiments. The BrdU label-chase experiments presented in this study thus allow us to quickly and easily demonstrate that the esophageal epithelia in rats and mice as well as D3-organoids contained the slow cycling/quiescent basal stem cell population (QBCs) that accounted around 4∼7% of total basal cells in the basal layers. These QBCs/stem cells were spatially and randomly located in the esophageal epithelial basal layer.

Isolation of label (BrdU+) and unlabel (BrdU-) subpopulation from label-chase experiments in rat with omic analyses demonstrated that hierarchical clustering in the methylation sites from WGBS could clearly separate QBCs from PBCs and SPBCs, indicating that although QBCs were spatially and randomly colocalized with PBCs in the basal layer, QBCs were unique in terms of epigenetic regulations at DNA methylation levels. These data were in agreements with previous studies showing that stem cells and differentiation progeny cells had distinct epigenomic landscapes. DNA Methylation would be one of the important epigenomic modifications for the stem cell maintenance, differentiation and reprogramming[44].

In contrast to scRNA-seq using esophageal tissues reported recently, which contained epithelial cells and other types of cells [25, 45], our scRNA-seq using D3-organoids which were composed of only esophageal squamous epithelial cells enabled us to determine squamous epithelial subpopulations in detail. These results together with in vivo DNA methylation profiling indicated that QBCs represented the stem cells with high levels of HDs and low levels of Wnt signaling in the esophageal stratified squamous epithelia. Among them, QBCs expressed not only the basal cell markers but also low levels of cell cycle markers, demonstrating that QBCs represented a group of cells with high levels of HD components (*Itgα6*, *Itgβ4*, *Col17a1*, *Dst* and *Plec*) and Wnt pathway negative regulators *(Senp2 and Prickle1)* in the esophageal stratified squamous epithelia. Pseudo-time cell trajectory showed that QBCs in the basal layer produced proliferating and/or differentiating cells in the basal layer, which, in turn, progressed into differentiating cells in the suprabasal layer and ultimately transforming into differentiated keratinocytes in the differentiated layer. Thus, these results indicated that QBCs represented the stem cells in the esophageal stratified squamous epithelia.

HD components, ITGα6 and ITGβ4, were reported as markers of esophageal stem cells in previous studies [17, 21, 23]. It was shown that isolated keratinocytes with SOX2^+^ITGβ4^hi^ITGα6^hi^CD73^low^ from the esophagi of mice could form more and larger organoids when compared with cells with ITGβ4^low^ITGα6^low^ [21], Using 3D organotypic sphere culture system, cells isolated from mice with CD49f^+^ (also known as ITGα6) CD24^low^CD71^low^ were shown to enrich esophageal stem cells that could display higher sphere-forming capacity and give rise to differentiated suprabasal cells[23]. Recently, other HD components, such as COL17A1, were also identified as esophageal epithelial stem cell maker by other study where the authors showed that stem cells in the human epithelium expressed high levels of COL17A1[25]. Moreover, studies showed that high expression of COL17A1 in mouse skin marked epidermal stem cells. In that study, expression levels of COL17A1 controlled stem cell competition and orchestrated skin homeostasis and aging[35]. Our results presented here were not only consistent with these data but also further demonstrated that perturbation HD components, COL17A1 and PLEC, inhibited esophageal keratinocyte organoid formation, morphogenesis and proliferation-differentiation homeostasis. Taken together, the results indicated that HDs were not only a prominent marker for the stem cells but also involved in regulating the stem cell maintenance and proliferation-differentiation homeostasis in the basal layer of the esophagus.

Our results also revealed that low levels of Wnt signaling had crucial roles in stem cell maintenance and proliferation-differentiation homeostasis in the mammalian esophageal epithelium. Wnt pathway were tightly linked with the stem cell maintenance and differentiation in multiple mammalian tissues [46, 47]. In intestinal niches, a gradient of Wnt signaling activities were found along the colonic crypt axis with the highest levels at the crypt bottom to maintain LGR5+ stem cells [48–52]. LGR5+ cells, resident at position +4 in a niche, had the lowest level of Wnt signaling than other LGR5+ cells in the niche, thus representing a long-lived, slow cycling/quiescent stem cell population[29, 53, 54]. As our omic results obtained from the scRNA-seq and WGBS pointed to QBCs that were enriched high expressions of Wnt signaling negative regulators, we manipulated Wnt signaling in D3-organoids, demonstrating that QBCs of esophageal epithelium were required low levels of Wnt signaling for their maintenances.

We explored the relationship between HD and Wnt signaling and found that not only high levels of HDs and low levels of Wnt signaling but also their cross-talk(s) and/or interplay(s) defined QBCs of the basal layer although the precise underlying mechanism(s) would be required for further investigations. Based on these studies, we propose that high levels of HDs (numbers) and low levels of Wnt signaling controlled, at least in part, by their components/regulators’ expression via epigenetic regulation at DNA level (DNA methylation) and their cross-talks/interplays at the basal cells define the stem cells, which are quired for self-renewal, maintenance and proliferation-differentiation homeostasis in the mammalian esophagus.

## Materials and methods

### Cell line and culture conditions

Human telomere reverse transcriptase (h-tert) immortalized rat normal esophageal epithelial cell line (RNE-D3, D3 for short) was established and preserved by our laboratory. D3 was cultured in DMEM/F12 (3:1) medium supplemented with 10% fetal bovine serum (Thermo Fisher Scientific), 8 ng/mL Cholera Toxin (CELL technologies), 5 ng/mL insulin (CELL technologies), 25 ng/mL hydrocortisone (CELL technologies), 0.1 ng/mL EGF (PeproTech) and 10 μM Y27632 (Topscience) in a humidified 37°C incubator supplemented with 5% CO_2_. D3-sh*Col17a1* cell lines were constructed by lentivirus transduction using following sequences: shRNA-1: 5’-GGACCTATCACAACAACATAG-3’.shRNA-2: 5’-GCAGACACATTCTCAACTATA-3’. D3-sh*PLEC* cell lines were constructed by lentivirus transduction using following sequences: shRNA-1: 5’-GCACAAGCCCATGCTCATAGACGAATCTATGAGCATGGGCTTGTGC-3’. shRNA-2: 5’-GCGCATTGTGAGCAAGCTACACGAATGTAGCTTGCTCACAATGCGC-3’.

### Organoid culture and in vitro BrdU-labeling

Generation of organoids was performed as described previously[21]. Briefly, D3 cells, D3-shCOL17A1 cells or D3-shPLEC cells were trypsinized into single cells and resuspended by ice-cold Matrigel (Corning). A droplet of 50 μL cell-Matrigel mixture was seeded into the bottom central of flat 24-well plates. After solidification in incubator, 500 μL medium of advanced DMEM/F12 (Thermo Fisher Scientific) supplemented with 1×penicillin-streptomycin (Thermo Fisher Scientific), 1×N2 supplement (Thermo Fisher Scientific), 1×B27 supplements (Thermo Fisher Scientific), 10 mM HEPES Buffer (CELL technologies), 1×GlutaMAX™ (Thermo Fisher Scientific), 1 mM N-acetyl-L-cysteine (Sigma Aldrich), 100 ng/mL recombinant murine EGF (PeproTech), 100 ng/mL recombinant murine Noggin (PeproTech), 100 ng/mL recombinant human R-Spondin1 (R&D Systems) and 10 μM Y27632 (Topscience) were added to each well. Organoids were grown for 10-12 days at 37°C in a CO_2_ incubator with the medium changed every other day. The organoid formation rate (OFR) was determined by calculating the percentage of the average numbers of organoids to the cell number initially seeded per well.

For BrdU labeling assay, 200 μM BrdU (Sigma-Aldrich) was added at 6th to 9th day of organoid culture. After 10 days of culture, organoids were collected from Matrigel by cell recovery solution (Corning) digestion. For subsequent BrdU immunofluorescence staining, the collected organoids were fixed in neutral fixative solution and embedded in OCT for following experiment. For Wnt inhibition experiment, Wnt inhibitor IWP-2(TOCRIS) and Srfp-2(R&D Systems) were added with BrdU at the same time on the 6th day of D3-organoid culture or D3-shPLEC-organoids culture and organoids were collected on the day 10. Another experiment with Wnt activator was that CHIR-99021(TOCRIS) was added at the beginning of D3-organoid culture and organoids collection was on day5, followed by H&E staining.

### Animals and in vivo BrdU-labeling

All procedures involving animals were carried out in accordance with the standards approved by ethical committee of National Cancer Center/National Clinical Research Center for Cancer/Cancer Hospital, Chinese Academy of Medical Sciences. Spargue-Dawley (SD) rats and BLAB/C mice, 4-5 weeks old, were purchased from BEIJING HUAFUKANG BIOSCIENCE COMPANY. Rats and mice were housed two and five per cage under standard conditions (24°C ±2°C, 20 relative humidity, 12-h light/dark cycles) and given access to standard rodent maintenance feed (Keao Xieli Feed, Beijing, China) and water ad libitum. Hygienic conditions were maintained by weekly cage changes. After completion of experiments, we sacrificed mice and rats by inhalation of anaesthetics with CO_2_.

The animals were intraperitoneally injected with BrdU of 100 mg/kg body weight once per 6 hours for 4 days and sacrificed at designed time points. The entire esophagus was obtained and fixed by 10% formalin overnight, followed by routine histological processing of H&E and immunohistochemistry staining. For immunofluorescence, the entire esophagus was fixed by 10% formalin overnight and dehydrated in 30% sucrose solution. Then the segmented esophagus was embedded in OCT compound and immediately frozen by liquid nitrogen to proceed with cryosection.

### Protein extraction and immunoblotting analysis

Cell samples were collected and extracted to obtain proteins according to the manufacturer’s requirements. Immunoblotting was performed as described previously [55]. The primary antibodies and dilutions were used against COL17A1(Abcam, 1:1000), PLEC (Abcam, 1:1000) and β-actin (Sigma Aldrich, 1:5000), respectively.

### Immunofluorescence

The OCT embedded esophagus tissue or organoids were cut into 5-10 µm sections using cryosection system. The sections were blocked with 10% normal goat serum (containing 0.2% TritonX-100) for 1 hour followed by incubation with primary antibodies against BrdU (Abcam, 1:200), BMI1 (Sigma, 1:200), OCT4 (Abcam, 1:200), SOX2 (Abcam, 1:500), Cytokeratin14 (Abcam, 1:1000), Cytokeratin13 (SantaCruz, 1:500), PCNA (CST, 1:2400), P63 (Abcam, 1:100), Integrin6 (SantaCruz, 1:500), Integrinβ4 (Abcam, 1:100), CD34 (Abcam, 1:100), P75 (Abcam, 1:50), DST (Affinity Biosciences, 1:200) and COL17A1 (Abcam, 1:200), respectively. Sections were counterstained with DAPI and sealed with Slowfade Diamond Antifade Mounted solution (ThermoFisher Scientific) for microscopic observation. The image fluorescence intensity was measured by Fiji Image J. All the images were converted to 8 Bit grayscale for plot profile analysis.

### Immunohistochemistry

Paraffin-embedded sections (5-10 µm) were deparaffinized and hydrated. Citrate buffer solution (PH 6.0) was used for microwave antigen retrieval for 10 minutes. Endogenous peroxidase was blocked by 3% hydrogen peroxide solution for 20 minutes. The sections were subsequently blocked with 10% goat serum for 1 hour and incubation with primary antibody at 4°C overnight, followed by HRP polymer incubation for 1 hour at room temperature. DAB solution was used for chromogenic reaction under microscopic observation. Then the sections were counterstained with Hematoxylin and sealed with neutral balsam for microscopic observation. The primary antibodies used were BrdU (SantaCruz, 1:200), Cytokeratin14 (Abcam, 1:5000), Cytokeratin13 (SantaCruz, 1:500) and PCNA (CST, 1:4000).

### FACS and Cell cycle analysis

Epithelial cells were obtained from rat esophagus as described previously[21]. Then these cells were fixed with 2% paraformaldehyde and then washed by PBS twice. Cells were penetrated with PBS containing 0.1% TritonX-100 for 5 minutes on ice and washed by PBS for 3 times. DNase I (TaKaRa) was added to each sample, incubating for 30 minutes at 37°C in the dark. After Washing, cells were stained with primary antibody anti-rat BrdU (SantaCruz, 1:200) and anti-rabbit CK14 (Abcam, 1:5000) for 30 minutes at room temperature. Subsequently, cells were incubated with goat anti-rat Alexa flour® 488 and donkey anti-rabbit APC (IgG H&L) for 30 mins at room temperature. Then the BD Flow Sorter was used to sort the BrdU+CK14+, BrdU-CK14+ and BrdU+CK14-cells. For cell-cycle assay, the final cell pellet was suspended in 400 μl of PBS containing a 1:1000 dilution of propidium iodide (PI) for 30 minutes at 37°C protected from light followed by flow cytometry examination. Obtained data were further analyzed by Flowjo software (version 10).

### q-PCR

For q-PCR, total RNA was extracted using the Trizol reagent (Ambion, USA) and reverse-transcribed to complementary DNA using the PrimeScript™ RT Reagent Kit (Takara, Dalian, China). Q-PCR was carried out using the SYRB Premix Ex Taq™ Perfect Real-Time system (Takara). The expression levels were normalized to that of the housekeeping gene GADPH. The primers were used following sequences: GAPDH_F: 5’- CATGCCGCCTGGAGAAAC -3’; GAPDH_R:5’- CCCAGGATGCCCTTTAGT -3’; Axin2_F:5’- GACAGCGAGTTATCCAGCGA -3’; Axin2_R:5’-GTGGGTTCTCGGGAAGTGAG -3’; Dvl1_F: 5’- ATGAGGAGGACAACACGAGC -3’; Dvl1_R:5’- AAGTGGTGCCTCTCCATGTT-3’.

### MethyIC-seq library construction, sequencing and data analysis

Samples (BrdU+CK14 +, BrdU-CK14+ and BrdU+CK14- cells) were isolated from the esophagi of rats labeled with BrdU for 4 days as described above. Extracted DNA samples would first be examined for concentration and purity to exclude degradation or contamination. Acegen Bisulfite-Seq Library Prep Kit (Acegen, Cat. No. AG0311) was applied for Whole genome bisulfite sequencing libraries construction according to the manufacturer’s instruction. In brief, 1 ng unmethylated Lambda DNA was mixed with 1 μg extracted genomic DNA, followed by sonication into approximately 200-500 bp fragments. Then the subsequent end-repaired, 5’-phosphorylation, 3’-dA-tailing and 5-methylcytosine-modified adapter ligation were performed. After being processed by bisulfite, PCR was operated for 10 cycles to amplify the DNA using Illumina 8-bp dual index primers. The constructed WGBS libraries were then analyzed by Agilent 2100 Bioanalyzer and finally sequenced on Illumina platforms using a 150×2 paired-end sequencing protocol. Agilent 2100 Bioanalyzer and qPCR was used for analyzing and qualifying the libraries. Illumina HiSeq X10 platforms with a 150×2 paired-end sequencing method was used for final sequencing.

The FastQC software (version 0.11.7) was used for quality control of the raw data, and the Trimmomatic software (version 0.36) was used for removal of adapters and unqualified data. The optimized data was mapped to the Rnor_6.0 Rattus reference genome using the BSMAP software (version 2.73). Data available for further analysis should comply with this standard: unique aligned reads, methylated cytosines with sequence depth coverage ≥5. Calculation of individual cytosine methylation level was performed using the ratio of sequenced CpG methylated cytosine depth to total CpG cytosine depth. Differentially Methylated Region (DMR) was established using Metilene software (version 0.2-7), defining ≥5 cytosine sites in the candidate region no more than 200bp distance to the neighboring cytosine (30bp for CHH). Average methylation levels differences of CG-DMR, CHG-DMR and CHH-DMR regions were all need to >0.1 between different populations. Finally, regions established as final DMRs should be in accordance with the following principles: 2D KS-test p-value <0.05, BH (Benjamini & Hochberg) corrected p-value < 0.05. To investigate biological process differences involved in DMR-related genes, Gene Ontology (GO) enrichment was performed (Q ≤ 0.05 was considered significantly enriched). Next, annotated genes with DMR overlapped on their gene body or upstream and downstream in 2kb were enriched for Kyoto Encyclopedia of Genes and Genomes (KEGG) functional analysis.

### Single-cell RNA sequencing and data processing

D3 organoids were collected from 24-well plate by Cell Recovery Solution digesting on ice for 2 hours. The deposited organoids were scattered by mixture digesting solution (containing 1× collagenase I, 1× collagenase IV, and 1× trypsin), followed by digestion for 30 minutes at 37°C. Then the treated organoids were centrifuged and resuspended as single-cell solution by PBS containing 0.04% BSA for further sequencing.

Resuspended single cells were embedded into a single-cell gel beads on a Chromium single cell controller (10× Genomics) with the application of single cell 3 ’Library and Gel Bead Kit V3 (10× Genomics, 1000075) and Chromium Single Cell B Chip Kit (10× Genomics, 1000074) following the manufacturer’s instructions. The wrapped beads containing individual cells, specific barcodes, UMIs (unique molecular identifiers), cell lysis solution and mixture needed for reverse transcription. After reverse transcription, obtained cDNA with specific barcodes and UMIs were mixed together for single-cell RNA-seq library construction using Single Cell 3’ Library and Gel Bead Kit V3. Then the final sequencing was operated using an Illumina Novaseq6000 sequencer with a sequencing depth of at least 100,000 reads per cell with pair-end 150 bp (PE150) reading strategy (performed by CapitalBio Technology, Beijing). FastQC software (version 0.11.2) was used for quality control, and the obtained data was mapped to the Rnor_6.0 Rattus norvegicus reference genome using Cell Ranger software (version 4.0.0). Barcode counting, UMI counting, and cell filtering were performed to achieve feature-barcode matrix and determine clusters using Cell Ranger software. Exclusion criteria for abnormal cell when gene number was less than 200, or gene number ranked in the top 1%, or mitochondrial gene ratio was more than 25%. After UMI normalization, PCA (Principal Component Analysis) and ten principal components were used to perform dimension reduction by K-means algorithm (version 0.17) and graph-based algorithm (version 0.17), respectively. Visualization was realized by t-SNE dimension reduction analysis (version 0.15). Then the enrichment analysis was performed using the top 20 marker genes of each cluster by means of KEGG and GO analysis (KOBAS software). Single-cell trajectories determined as pseudotime were built with Monocle (version 2.4.0). The WGCNA R software package (version 1.51) was used for weighted correlation network analysis. Sub-clusters would be generated from every defined cluster according to above clustering result, expression of genes will be further calculated, and the relative expression level of specific genes were presented as violin plots. Gene Set Enrichment Analysis (GSEA) was processed by GSEA software (version 2.2.2.4), which uses predefined gene sets from the Molecular Signatures Database (MSigDB version 6.2). To further verify the accuracy of cell cluster definition, GSVA (Gene Set Variation Analysis) scores for given biological process (including fatty acid metabolism, G2/M cell cycle, glycolysis, oxidation phosphorylation) and NRF2-regulated redox state were calculated in each cell cluster using GSVA software (version 1.30.0). NRF2 regulated gene set including 469 genes were downloaded from GSEA website (http://www.gsea-msigdb.org/gsea/msigdb/genesets.jsp?letter=N).

### Statistical analysis

Student’s t-test and Two-way ANOVA were performed for analyzing statistic differences between groups, and p<0.05 was considered of significance. Data were presented as Mean ± SD. GraphPad Prism 7.0 was used for analysis.

### Study approval

All experiments involving animals were complied with the standards approved by ethical committee of National Cancer Center/National Clinical Research Center for Cancer/Cancer Hospital, Chinese Academy of Medical Sciences.

## Acknowledgments

This work was supported by the National Natural Science Foundation of China (NSFC: 81972572 to WJ), and Chinese Academy of Medical Sciences (CAMS) Innovation Fund for Medical Sciences (CIFMS) (Grant no.2016-I2M-1-001 to WJ and S-HL). During this study, Professor Shih-Hsin Lu passed away in 2019. We all miss him.

## Author contributions

YY and GD contributed to study design, experiment operation, data interpretation and manuscript writing. LQ, YH, YX and LX contributed experiment operation and data interpretation: S-HL, WJ and XY contributed project supervision and data interpretation. WJ and XY contributed manuscript writing. All authors reviewed and approved the final manuscript.

## Supporting information

**Figure S1.**
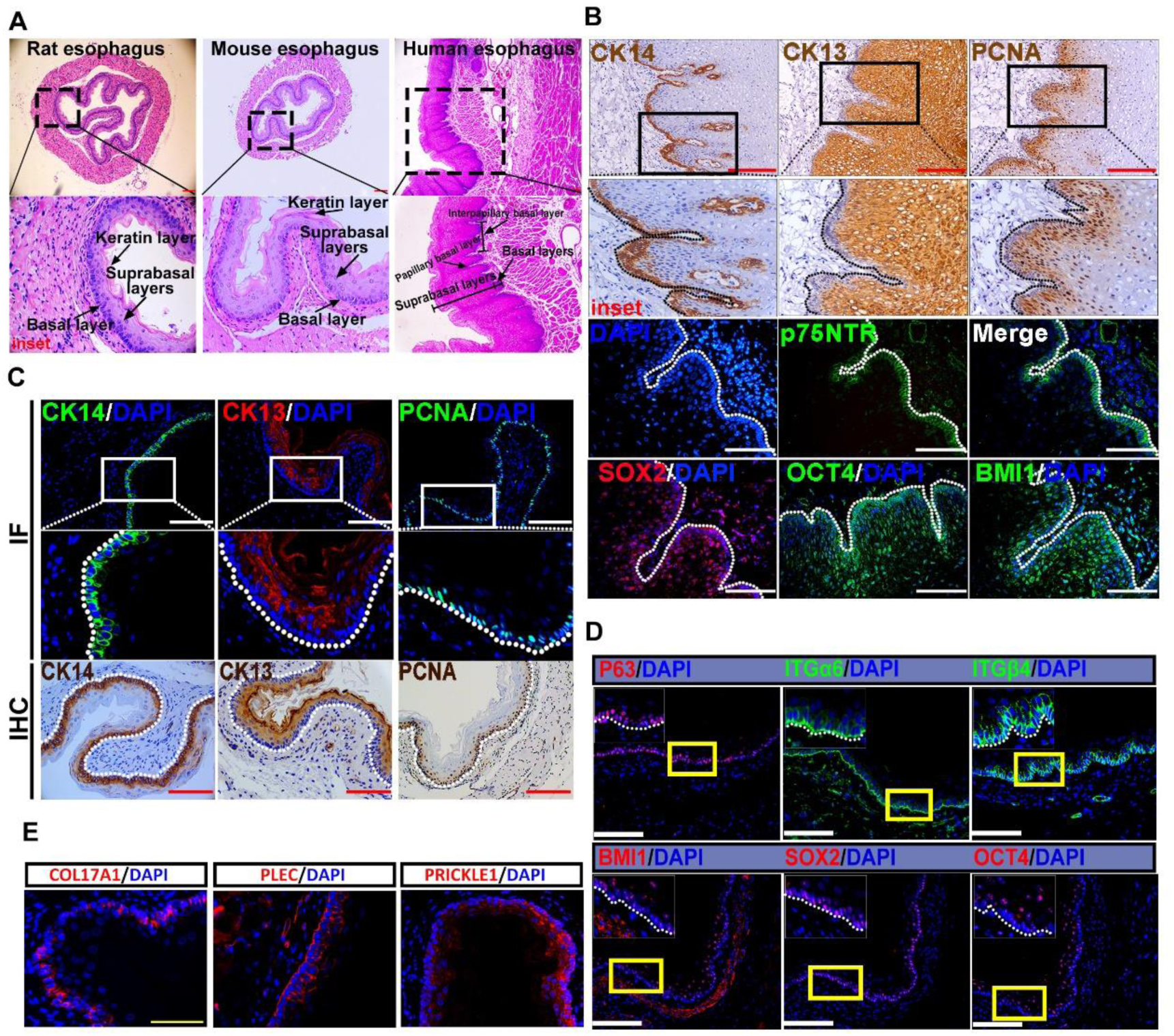
Characterization of rodent and human esophageal epithelium. (a) H&E staining of normal rodent and human esophagus cross-section. Rodent esophagus with endodermal structures including basal layer, suprabasal layers and keratin layer. Human esophagus with endodermal structures including suprabaseal layers, papillary and interpapillary basal layer. Immunostaining of CK14, CK13, PCNA, P75, SOX2, OCT4 and BMI1 of human esophageal sections. (C) Immunostaining of CK14, CK13, PCNA, (D) P63, ITGα6, ITGβ4, BMI1, SOX2, OCT4 and (E) COL17A1, PLEC and PRICKLE1 of rat esophageal tissue sections. Dotted line marks the basement membrane. Scale bars: 100μm.

**Figure S2.**
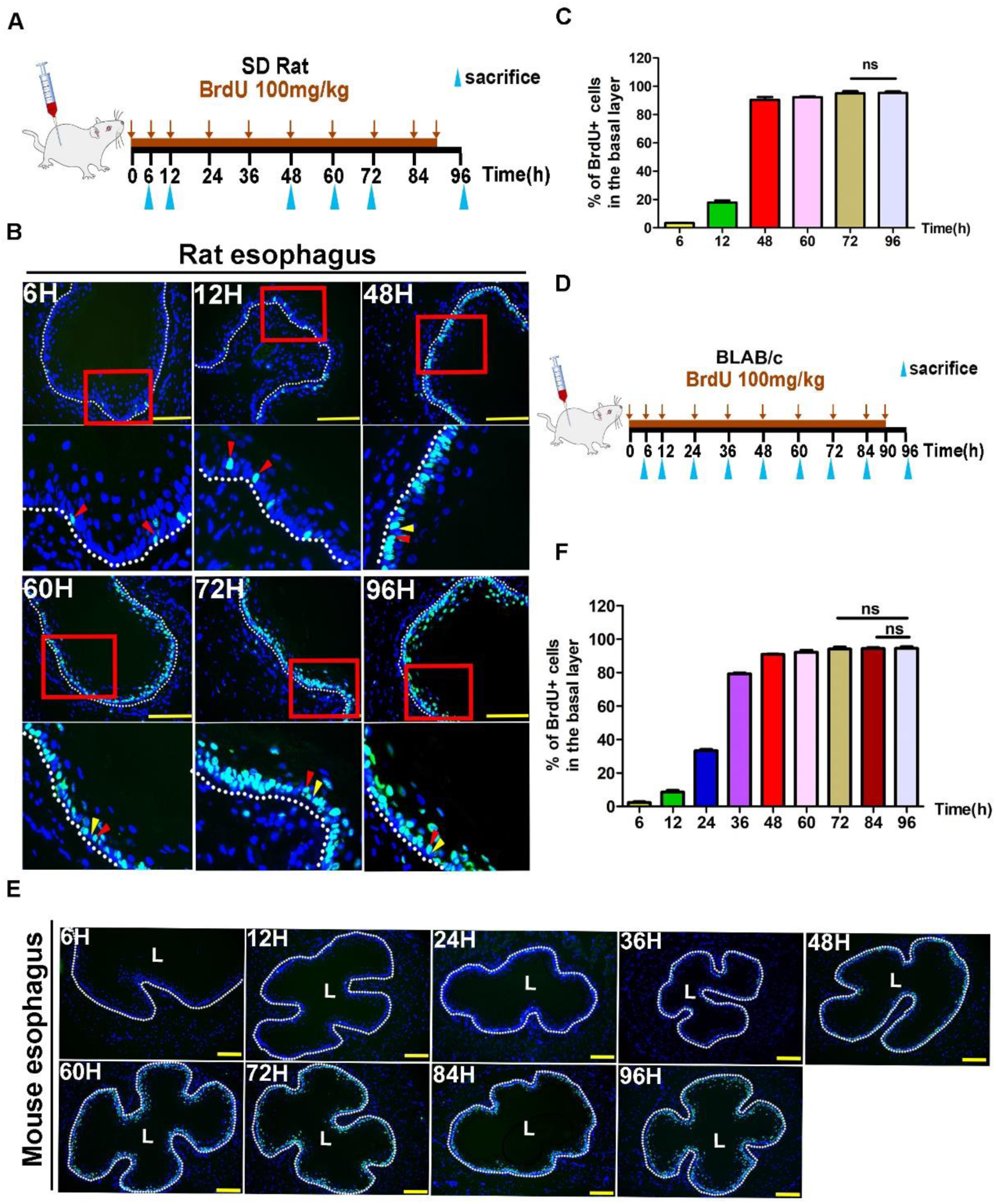
Rodent esophageal basal layer exists a small relatively slow cycling/quiescent cell population. (A) Schematic illustration of BrdU-labeling experiment of SD rats. SD rats were injected with BrdU of 100 mg/kg body weight once per 6 hours for 96 consecutive hours and sacrificed at five different time points. (B) Immunofluorescence staining of BrdU (green) of rat esophageal sections at listed time points counterstained with DAPI (blue). Red triangle arrows indicate BrdU+ cells; the yellows indicate BrdU- cells. (C) The percentage of BrdU+ cells in the basal layer of rat esophageal epithelium at listed time points. (n=5, n represents 5 intact basal layers of esophageal epithelium counted at each time point). (D) Schematic illustration of BrdU- labeling experiment of BLAB/C mice. BLAB/C mice were injected with BrdU of 100 mg/kg body weight once per 6 hours for 96 consecutive hours and sacrificed at nine different time points. (E) Immunofluorescence staining of BrdU (green) of mouse esophageal sections at listed time points counterstained with DAPI (blue). (F) The percentage of BrdU+ cells in the basal layer of mouse esophageal epithelium at listed time points. (n=5, n represents 5 intact basal layers of esophageal epithelium counted at each time point) Data are represented as the mean +/- SD for percent analysis (*p < 0.05, **p < 0.01, ***p< 0.001). “L” indicates the lumen; dotted line marks the basement membrane. Scale bars: 100μm.

**Figure S3.**
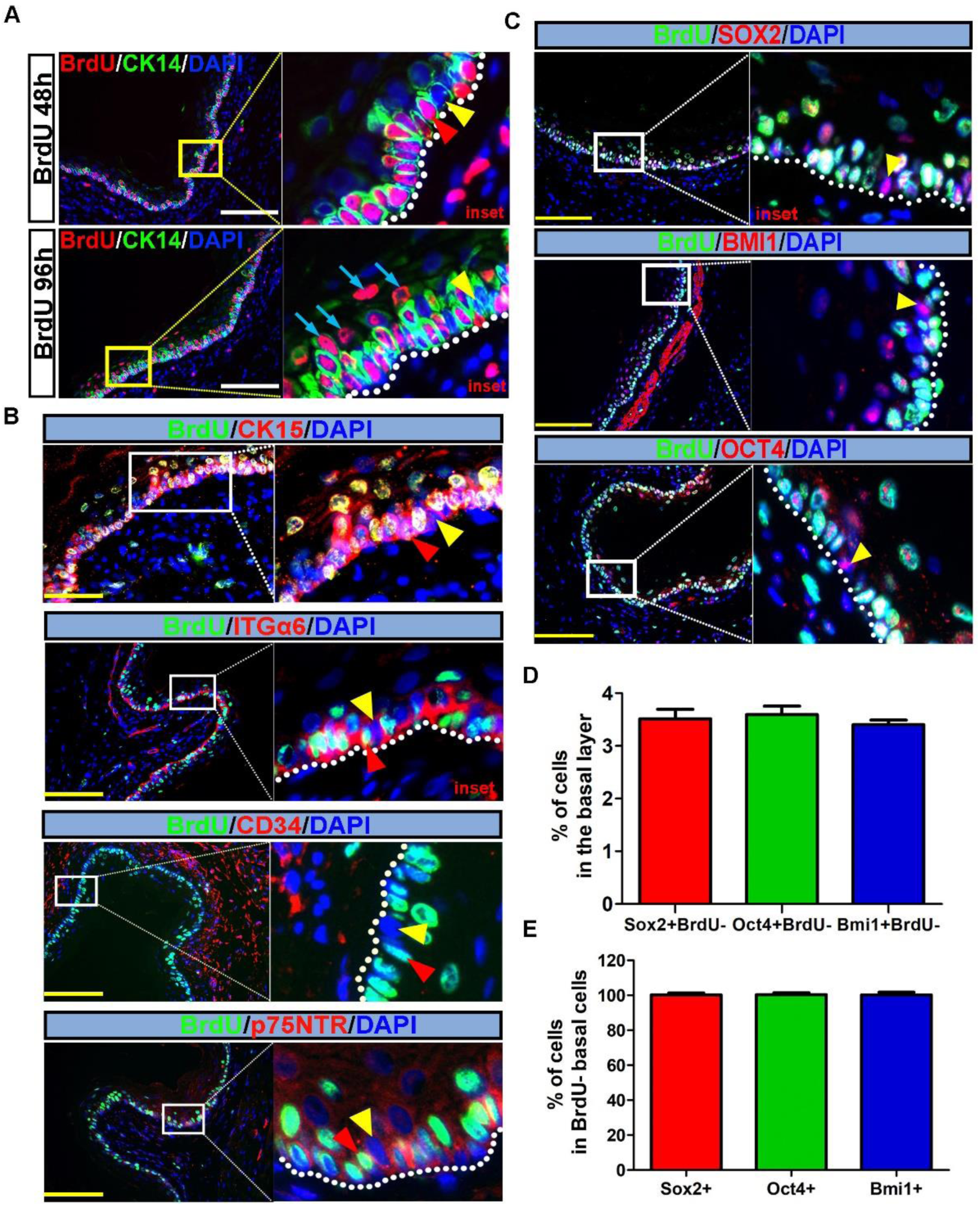
Rat esophageal slow cycling/quiescent basal cells co-immunostaining with stemness markers. (A) CK14 (green) and BrdU (red) co-immunostaining of rat primary esophageal tissue section counterstained with DAPI (blue). BrdU+ and BrdU- basal cells both expressed CK14 at BrdU- labelling 48 hours and 96 hours. (B) Colocalization of BrdU (green) with potential esophageal stemness markers CK15 (red), ITGα6 (red), CD34 (red) and P75NTR (red), respectively of the rat esophagus sections of BrdU 96h. (C) Colocalization of BrdU (green) with stemness-related markers SOX2 (red), BMI1 (red) and OCT4(red), respectively of the rat esophagus sections of BrdU 96h. (D) The percentage of SOX2+BrdU- cells, BMI1+BrdU- cells and OCT4+BrdU- cells in the basal layer cells that calculated of co-immunostaining were ∼4%, which were consistent with the percentage of BrdU- cells. (n=3). (E) The percentage of SOX2+ cells, BMI1+ cells and OCT4+ cells of BrdU- basal layer cells were almost 100% by manual counting (n=3). Insets panels represents magnification of regions of interest displayed by white or yellow rectangles. Yellow triangle arrows indicate BrdU- cells; the reds indicate BrdU+ cells. Dotted line marks the basement membrane. Scale bars: 100 μm. Data are represented as the mean +/- SD for percent analysis (*p < 0.05, **p < 0.01, ***p < 0.001).

**Figure S4.**
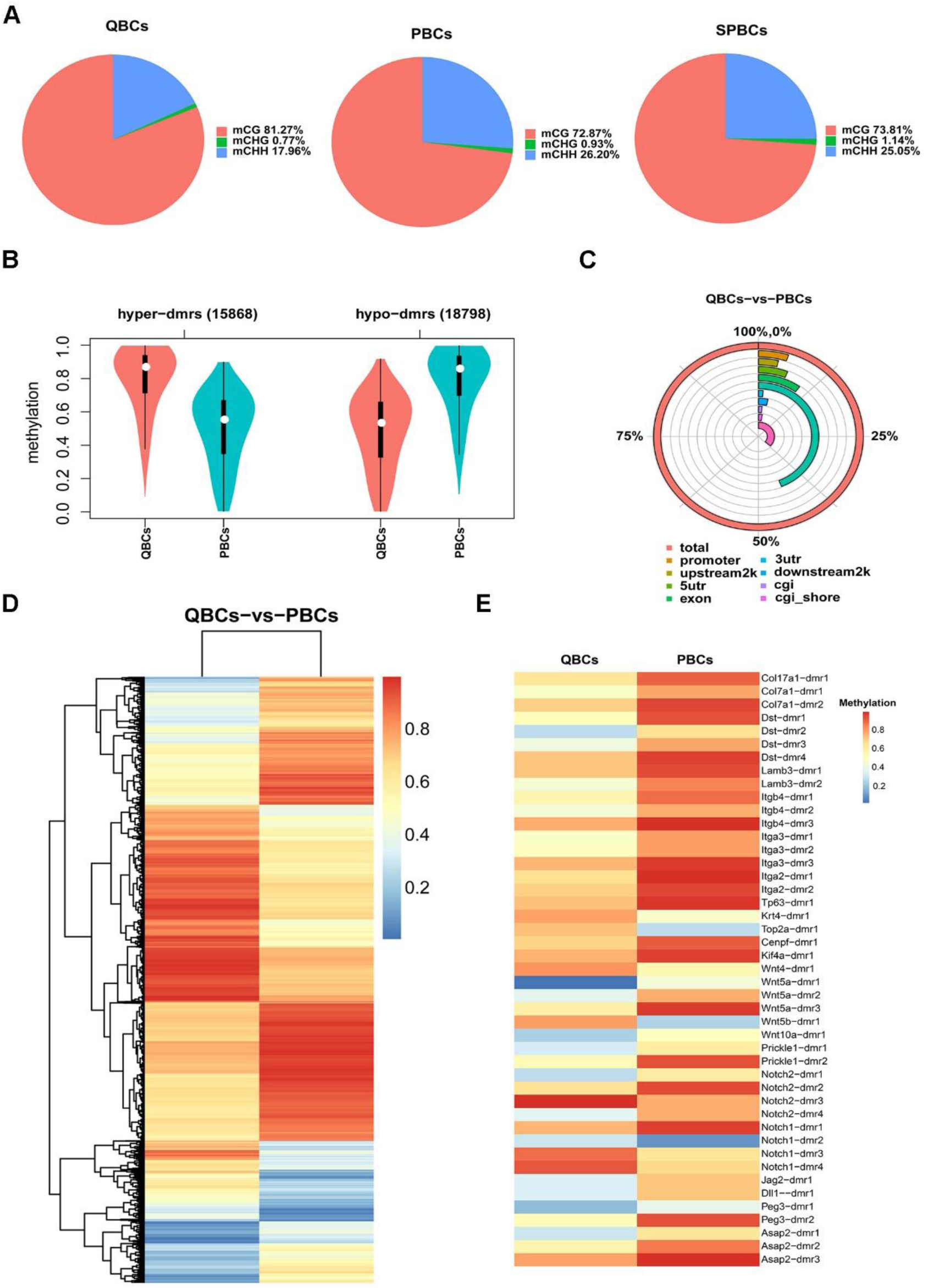
Differential methylation analysis among QBCs, PBCs and SPBCs. (A) Distribution map of methylated C sites of each population. Different colors represent methylated C sites in different contexts, and the size of each area represents the proportion of methylated C sites in the corresponding context. (B) DMR average methylation level distribution violin plot between QBCs and PBCs. DMR represent the differentially methylated regions. Hyper-dmrs represent the dmrs that are hypermethylated, and hypo-dmrs represent the dmrs that are hypomethylated. (C) Genomic functional element methylation map between QBCs and PBCs. (D) DMR methylation level clustering heat map between QBCs and PBCs. (E) DMR methylation level heat map of representative genes between QBCs and PBCs.

**Figure S5.**
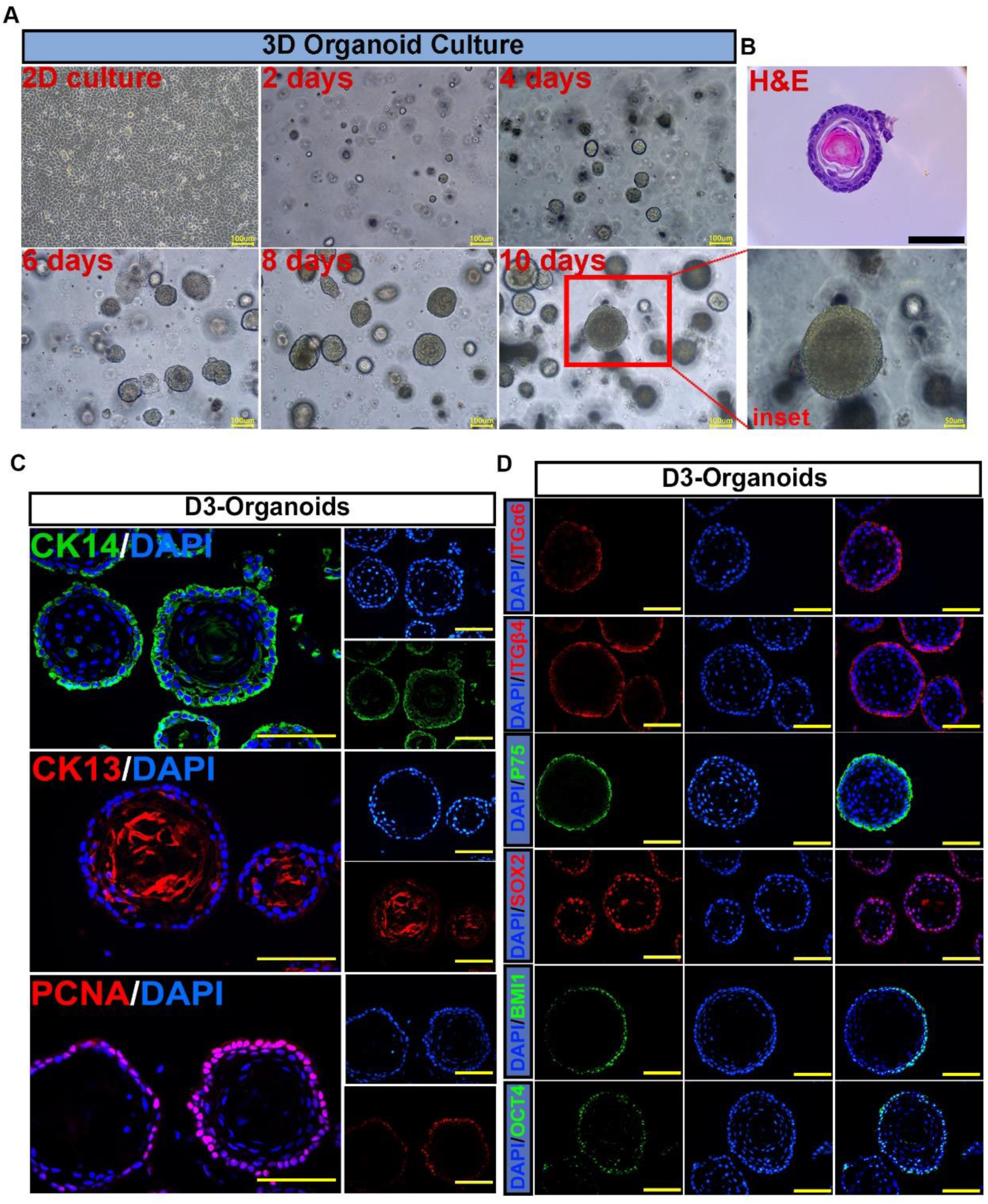
Characterization of rat esophageal organoids derived from immortalized normal rat esophageal keratinocyte cell line D3. (A) The expanding course of Immortalized normal rat esophageal keratinocyte cell line D3 cells with conditional culture to form normal and typical esophageal organoids. D3 cells in 2D culture were enzymatically dissociated and filtrated to prepare single-cell suspensions with Matrigel to initiate organoid culture in 10 days. Scale bars: 100μm. Inset showed the representative image of a normal and typical rat esophageal organoid derived from D3 cells (D3-organoid) in bright field. Scale bars: 50μm. (B) The representative image of H&E staining of D3-organoid. Scale bars: 100μm. (C) Immunofluorescence staining of CK14 (green), CK13 (red) and PCNA (red) counterstained with DAPI (blue) of D3-organoids. Scale bars: 100μm. (D) Immunofluorescence staining of esophageal stemness makers, ITGα6, ITGβ4, P75, SOX2, OCT4 and BMI1 of D3-organoids. Nucleus were counterstained with DAPI (blue). Scale bars: 100μm.

**Figure S6.**
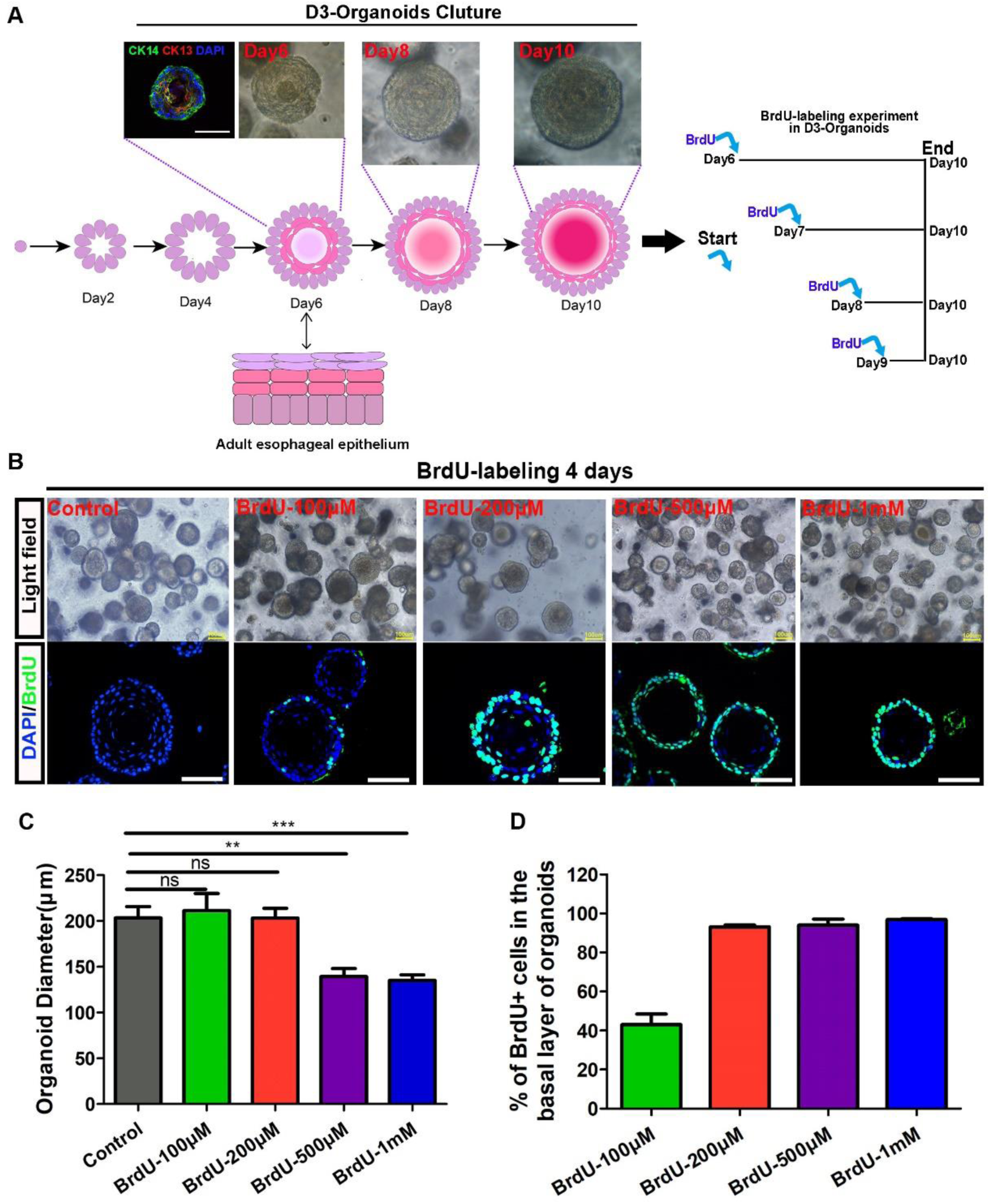
BrdU-labeling experiment of rat esophageal organoids derived from immortalized normal rat esophageal keratinocyte cell line D3. (A) Schematic illustration of BrdU-labeling experiment of rat esophageal organoids derived from D3. (B) The growth of D3-organoids was observed under different BrdU concentrations and immunofluorescence staining of BrdU was performed. (A) The diameters of D3-organoids were calculated under different BrdU concentrations. (n=5, n presents five random microscope fields,200x). (D) The percentage of BrdU+ cells in the basal layer of D3-organoids under different BrdU concentrations. (n=8, n presents eight random microscope fields,400x). Scale bars:100μm. Data are represented as the mean +/- SD for percent analysis (*p <0.05, **p < 0.01, ***p < 0.001).

**Figure S7.**
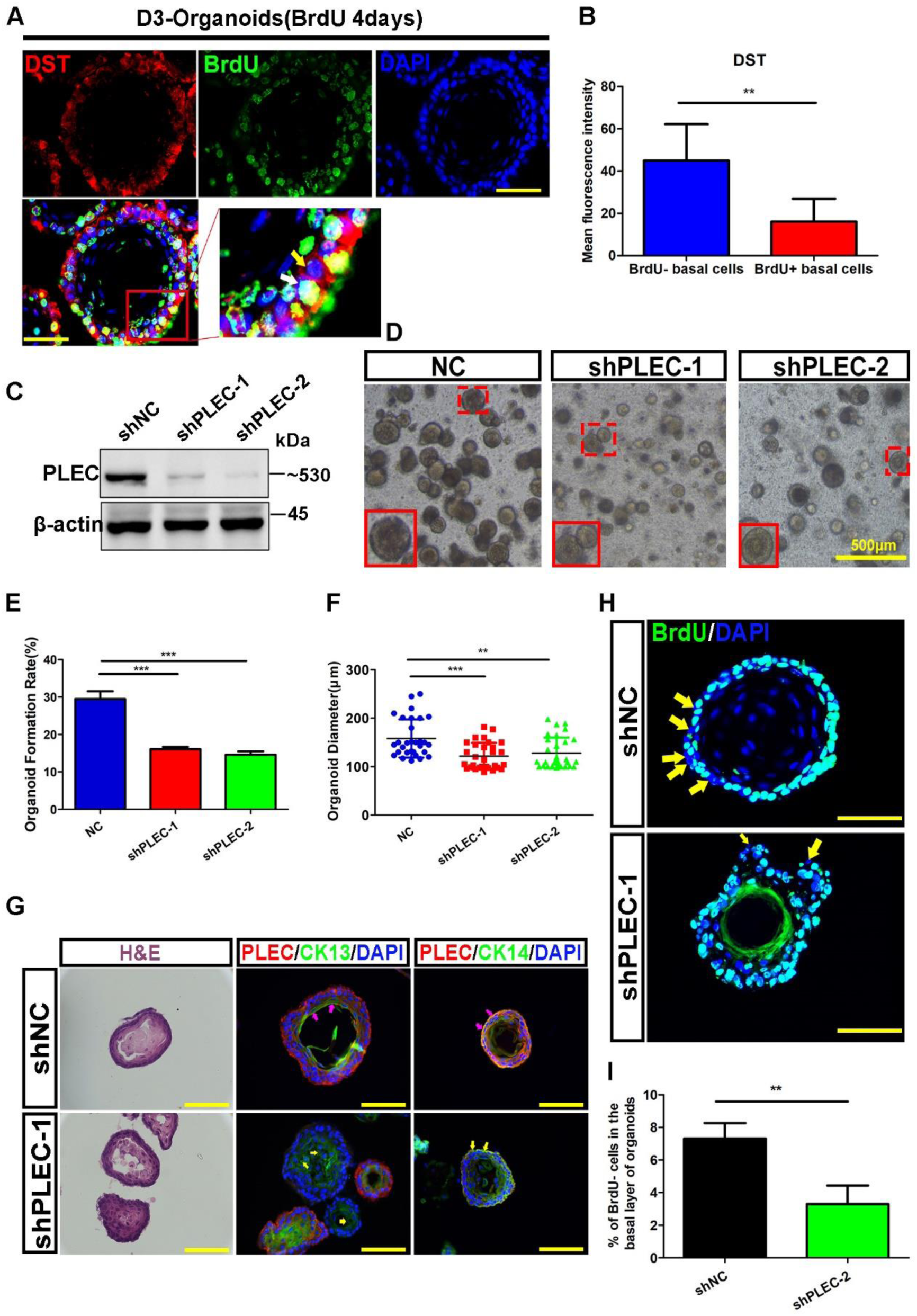
Hemidesmosome (HD) components in stem cell maintenance and proliferation-differentiation homeostasis of rat esophagus and organoids. (A) BrdU- basal cells had higher espression of DST expression than BrdU+ basal cells by immunofluorescence staining in D3 organoids. Scale bars:100μm. (B) The mean fluorescence intensity of DST expression was counted corresponds to (A). (C) Western Blotting verification of D3-sh*PLEC* cell line construction. (D) The representative images of light field of organoids at day 10. PLEC knockdown significantly inhibited organoid formation and growth. Scale bars: 500μm. (E) Quantification of organoid formation rate after PLEC knockdown. (F) Quantification of organoid diameter after PLEC knockdown. (G) H&E staining and immunofluorescence staining of intermediate filaments (CK13 and CK14) of organoids showed significant self-organization perturbation presented as uneven basal layers and abnormal distribution of CKs after PLEC knockdown. (H) Immunofluorescence staining of BrdU of D3- organoids labeled for 4 days after PLEC knockdown. The yellow arrows indicated the BrdU- cells. Scale bars: 100μm. (I)The percentage of BrdU- cells in the basal layer of D3-organoids after PLEC knockdown. (n=6, n presents six random microscope fields,200X). Data are represented as the mean +/- SD for percent analysis (*p < 0.05, **p < 0.01, ***p< 0.001).

**Figure S8.**
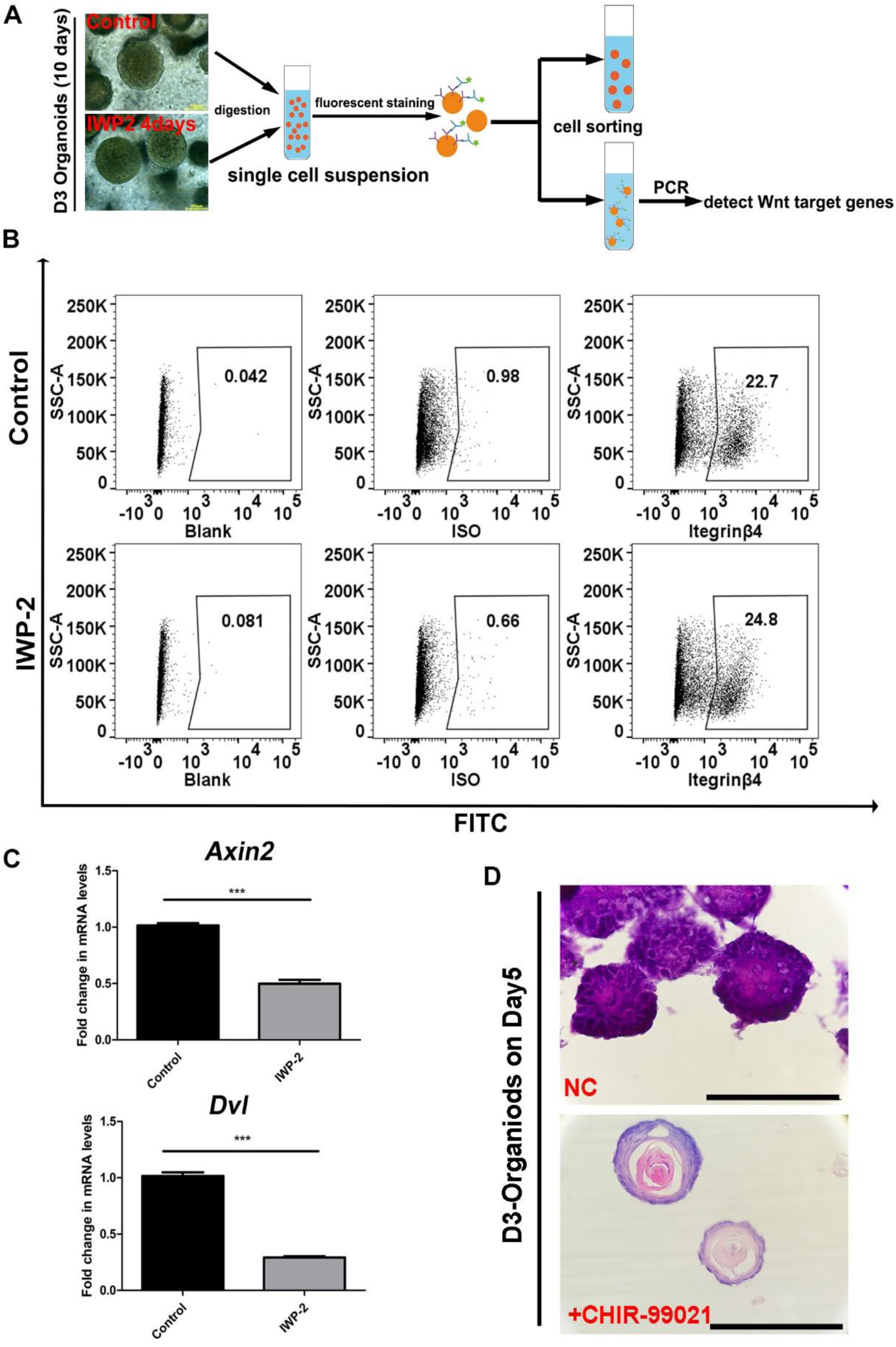
Wnt inhibition in rat esophageal organoids. (A) Experimental procedure for obtaining basal cells from organoids Scale bars: 100μm. (B) ITGβ4+ basal cells were sorted from D3- organoids by flow cytometry. (C) The mRNA expression of Wnt downstream target genes in the groups with Wnt inhibitor IWP-2 and control. (D) The D3-Organoids on day5 showed higher degree of differentiation treated with Wnt activator CHIR-99021. Scale bars:100μm.

